# Distinct roles of hippocampus and neocortex in symbolic compositional generalization

**DOI:** 10.1101/2025.08.19.671090

**Authors:** Zilu Liang, Leonie Glitz, Michelle Briana Hefner, Denis Lan, Miriam Klein-Flugge, Christopher Summerfield

**Author notes:** Corresponding authors (Z.L.), (M.K.F.), (C.S.). Senior authors (Miriam Klein-Flugge, Christopher Summerfield).

## Abstract

Humans can combine symbols to generate new meanings. Here, we studied the regional neural mechanisms that might make this possible. We asked participants to combine two discrete, symbolic features (a shape and a colour) to make a novel spatial inference. BOLD data suggested that the hippocampus encoded elementary visual attributes in a high-dimensional, parallel format that permitted flexible individuation. In ventromedial prefrontal cortex (vmPFC), posterior parietal cortex (PPC) and primary visual cortex (V1), neural patterns for novel stimuli (composites) could be predicted as a linear combination of signals for familiar stimuli (elements). In vmPFC, this composition occurred in a high-dimensional format, but in PPC and V1, it took place in a low-dimensional, spatial, response-consistent frame of reference. These data offer new insights into the neural circuit underlying compositional generalization.

## Introduction

The natural world is composed of discrete entities, such as words or objects. A major challenge for any cognitive system is to make inferences about entirely novel entities – those whose properties cannot be inferred by interpolation or extrapolation from extant knowledge distributions [1–4]. For example, European naturalists first arriving in Australia were baffled by a taxidermized *ornithorhynchus anatinus* (or duck-billed platypus), which they initially believed to be a hoax made by stitching together other animals. However, despite this scepticism, they might have predicted that this strange creature could swim (because it had a tail like a beaver) and that it laid eggs (because it had a beak like a bird). In other words, they could make inferences about an entirely novel entity by composing knowledge relevant to its elemental features. We and others have called this capacity ‘compositional generalization’ [5–12].

Across the 20^th^ century, compositional generalization was widely cited as the secret of human intelligence [13,14]. However, computational models of that era mostly ignored implementational questions about how composition occurs in the brain. Since then, neuroscientists have developed sophisticated theories of neural coding, proposing that stimuli are represented in neural population vectors, with the geometry of these vectors defining the space of possible computations [15,16]. For example, representations may be high-dimensional, making them highly separable in memory, or low-dimensional, supporting generalisability of knowledge [17–19]. Distinct variables (such as the identity and location of a stimulus) may be represented with orthogonal neural vectors, so that stimulus representations can be ‘factorised’ into elementary components, or with parallel neural vectors, which allow equivalences to be drawn between otherwise incommensurable features [5,20–23]. This perspective on neural coding thus provides a rich framework for understanding how composition occurs at the neural level.

Significant uncertainty remains, however, about how compositional generalization is implemented at the macroscopic level. Whilst a range of different brain regions have been implicated, it has been challenging to pinpoint their precise computational role. The ventromedial prefrontal cortex (vmPFC) and hippocampus (HPC), which are involved in the retrieval and integration of stored memories or task elements [24,25], are prime candidates. The hippocampus supports binding elements into relational structures, such as encoding the spatial and temporal organization of experiences [26], extracting graph-like transitions between events [27], and forming conjunctive representations of building blocks in specific relational configurations [7]. The vmPFC, on the other hand, plays a role in integrating individual components of a concept [9], such as performing semantic composition [28], representing complex rules composed from multiple single elements [29], inferring the value of novel options (such food items) by combining known components [30–32]. To do so, it may use an additive neural code, whereby neural patterns of elemental concepts are superimposed to form a compound concept (for instance, queen ≈ king - man + female) [33,34]. The precise brain regions involved, and the computations they may variously employ to produce new ideas from existing building blocks, thus remains an unsolved challenge in systems neuroscience.

Here, we studied the neural geometry underlying compositional generalization across the human brain. Whereas rodents and monkeys learn to compose very slowly (or not at all), humans can rapidly combine symbolic information to make novel inferences, even without instruction. We capitalised on a previously described task [8] in which participants viewed a cue composed of two discrete features (one of five animal shapes in one of five colours), and had to look for hidden treasure on a two-dimensional arena. The task required participants to learn that colour and shape mapped onto a distinct coordinate in the x- and the y-dimension of the arena respectively. However, during training they only ever received feedback on a subset of stimuli (an exhaustive set of colours for a single shape, and an exhaustive set of shapes for a single colour) so that successful test performance (with a novel combination of colour and shape) required them to compose knowledge of the relevant x- and y-position.

We recorded BOLD signals as a proxy for neural activity using fMRI, a brain imaging method with relatively poor spatial resolution, but which has been successfully used to probe neural geometry [35,36]. We posed three main questions. Firstly, we asked whether the neural code for colours and shapes would be high-dimensional (allowing individuation of colours and shapes) or low-dimensional (encoding the relative spatial position of the required response) during composition, and how the coding axes for the two features (shape and colour) were related to each other. Secondly, we hypothesised that neural activity elicited by (novel) test stimuli could be predicted from a linear combination of training stimulus activity, and asked whether this model fit the fMRI data, in either a naïve feature-based or distance-modulated fashion. Finally, using data from a separate localiser task, we asked whether composition occurred in a spatial or non-spatial frame of reference.

## Results

On day 1, participants learned the task over 6 cycles, each consisting of two training blocks followed by one test block (**Fig S1a**). On each trial they were shown a coloured (animal) shape and asked to predict where treasure would be found on a circular arena. They responded by moving a small pirate figure from a random position to the chosen location, using the arrow keys. Stimuli were associated with treasure locations organised on a 5 x 5 grid, with colour denoting one Cartesian coordinate (*x* or *y*) and shape the other. Coloured shapes on training trials (n = 9 per block) mapped to a single row and column in this array, and responses received fully supervised feedback (**Fig S1b**). Test trials (n = 16 per block), mapped onto remaining locations, but received no feedback (**Fig S1c**), obliging participants to use the knowledge learned during training to predict the correct response at test. Note that unlike in previous work [31], the cues were discrete and arbitrary (‘symbolic’) and thus participants could not learn to solve the compositional task with a simple rule. We call this task the ‘treasure hunt’ task.

Participants returned on day 2 and after a short refresher they entered the scanner and performed a version of the task in which training and test trials were intermixed, and no feedback was given (**Fig 1c**). We focus our analysis on the cue presentation period during which participants did not yet know which motor response would be required (because the pirate started in a random location when it appeared later in the trial, so participants could not plan their response trajectory).

**Figure 1.**
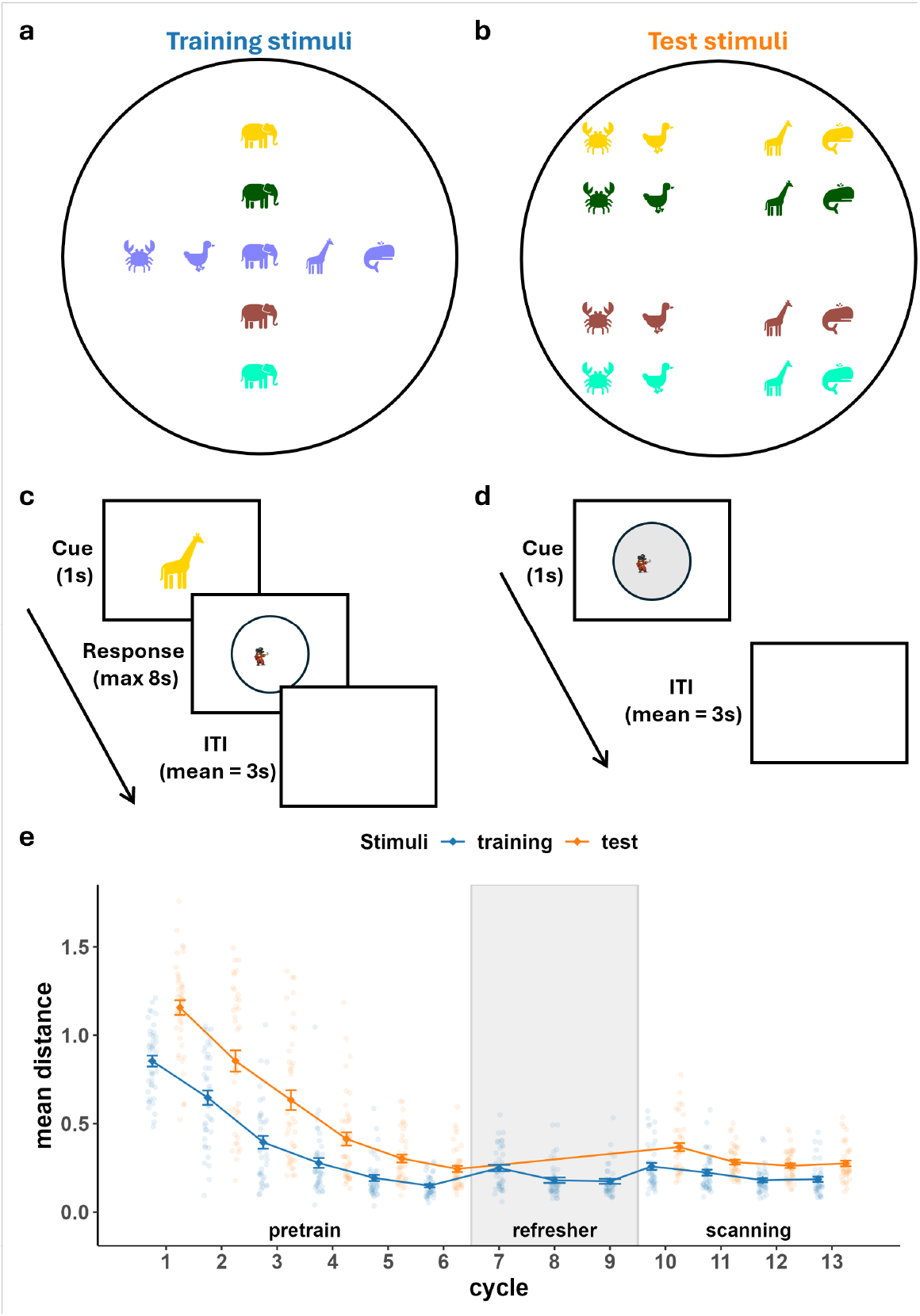
Task design and behaviour performance. (a-b) Example stimuli shown in their ground-truth locations on the map: (a) training stimuli (b) test stimuli. (c-d) Trial timelines of task performed in the scanner: (c) the treasure hunt task; (d) the spatial localiser task. (e) Performance of generalizers measured by distance to correct location. Translucent dots indicated individual generalizers. Solid lines indicated average performance of each participant group.

As observed previously [8], most participants successfully predicted the treasure locations for both training stimuli and test stimuli (‘generalizers’), but a subset of participants (‘non-generalizers’) either failed to learn the task or generalize according to the compositional rule we specified as ground truth. We classified participants as ‘generalizers’ (*n* = 41) or ‘non-generalizers’ (*n* = 15) using the same criterion as previously [8]. We plot the per-cycle performance for generalizers in **Fig. 1e** (see **Fig. S2** for a comparison between behavioural performance of non-generalizers and generalizers). We studied the neural representations of the 41 generalizers in three anatomically defined regions of interest (ROIs) that have been previously implicated in the learning of relational structure: the posterior parietal cortex (PPC) [37,38], the hippocampus (HPC) [39–42] and the ventromedial prefrontal cortex (vmPFC) [24,43,44]. We also included the primary visual cortex (V1), initially as a control, although we found that it also exhibited interesting neural geometry. These ROIs are shown in **Fig. 2a-d**.

**Figure 2.**
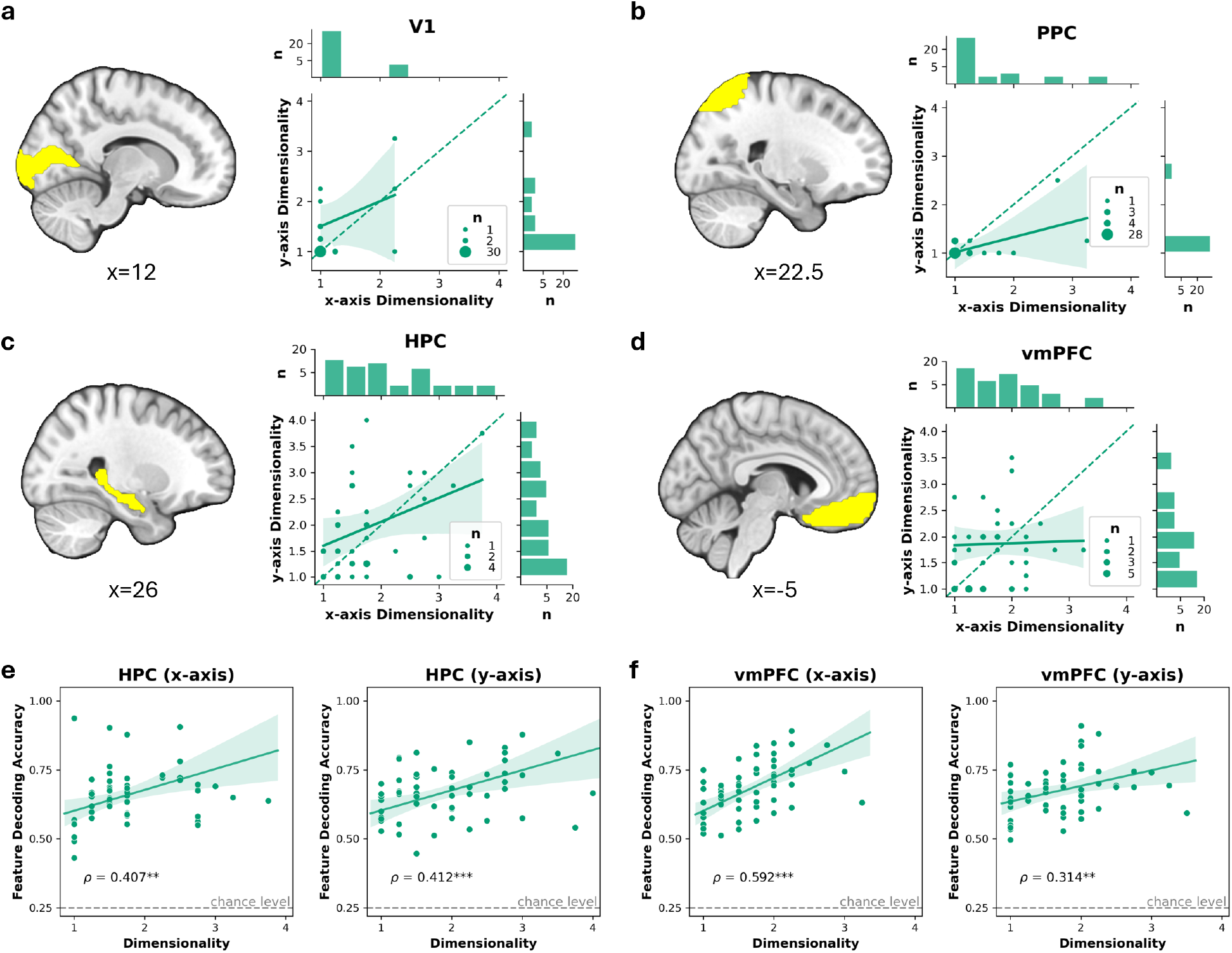
Dimensionality of training stimuli representation. (a-d) Dimensionality of x-axis and y-axis training stimuli representation in the four ROIs: early visual area (V1), posterior parietal cortex (PPC), hippocampus (HPC) and ventromedial prefrontal cortex (vmPFC). vmPFC and HPC showed higher dimensional codes than V1 and PPC. (e-f) Correlation between dimensionality of representation and feature decoding accuracy in HPC (e) and vmPFC (f). Asterisks indicate significance of the partial correlation coefficient against zero: ^***^ P<0.001, ^**^ P<0.01, ^*^ P<0.05.

### Dimensionality of neural codes during composition

To successfully locate the treasure for test stimuli, participants had to infer independent rules mapping discrete colours/shapes to the locations on the x- and y-axis. A maximally efficient encoding format for these rules would compress the discrete features into two one-dimensional neural codes (one for each axis). To test if any of the regions show such compressed code, we first estimated the dimensionality of the representations of individual rule from the training stimuli (**Fig. 3a**).

**Figure 3.**
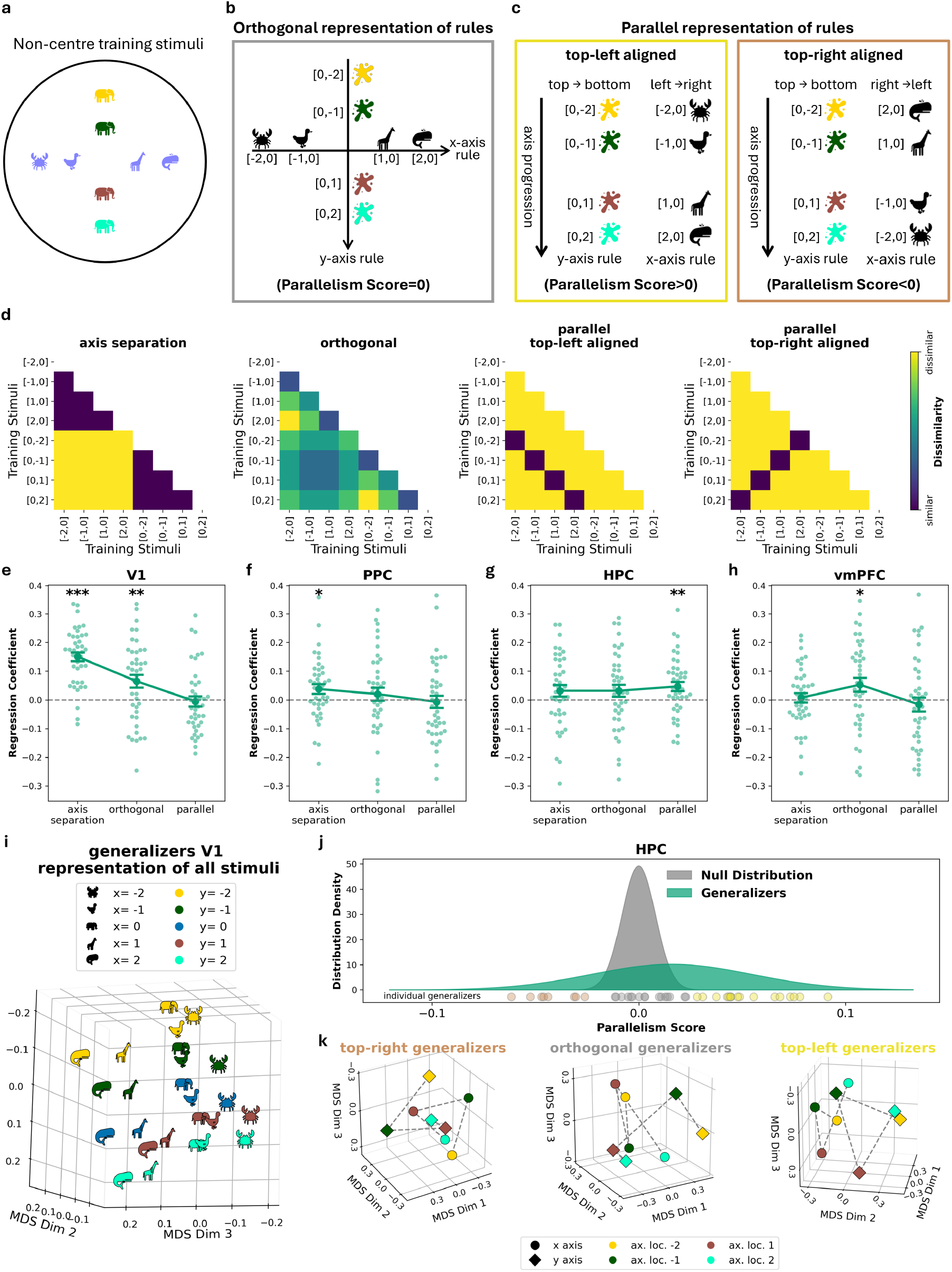
Parallel and orthogonal representation of training stimuli on different axes. (a) The non-centre training stimuli on x-axis and y-axis used in the analyses of dimensionality of individual rule representations and the geometric relationship between rules. (b-c) Schematic illustration of (b) an orthogonal representation of rules and (c) a parallel representation of rules. (d) Three model RDMs in the cross-validated RSA analysis. The axis separation model RDM assigned a distance of zero to all within-axis stimuli pairs and a distance of one to all between-axis stimuli pairs. The orthogonal model RDM corresponded the distance on the ground-truth map. The parallel RDMs assign a distance of zero to all stimuli pairs that had corresponding axis locations (depending on the alignment scheme between axes) and a distance of elsewhere. (e-h) Cross-validated RSA comparing training stimuli representation to three model RDMS shown in (d): axis-separation, orthogonal and parallel models. Dots represent individual generalizers. Error bars indicate mean ± standard error. Asterisks indicate results of permutation test against zero: ^***^ P<0.001, ^**^ P<0.01, ^*^ P<0.05. (h) Neural geometry of the generalizers’ representation of all stimuli in V1 revealed by multi-dimensional scaling (MDS). Note how it resembles the ground-truth map in **Fig. 1a-b**. (i) Observed and null distributions of parallelism scores (PS) in HPC in generalizers. Shaded areas show the normal distributions fitted to the observed data (in green) and the null (in grey). For visualization purpose, generalizers were classified into different axis-alignment schemes based on the relationship between their PS and the null. Coloured discs represent the observed PS of individual generalizers. Colours of the discs indicate the alignment scheme: top-right aligned (PS was significantly smaller than the null), orthogonal (PS did not deviate from the null), and top-left aligned (PS was significantly larger than the null). (j) The neural geometry of different scheme visualised using MDS. Training stimuli from the same training axis (indicated by shapes) are connected with dashed lines. Colours indicate the locations of stimuli on their respective axes. Note how lines linking the same colour pair in square and diamond were: parallel but inverted in the left panel, orthogonal in the middle panel, and parallel on the right panel.

We estimated the dimensionalities in each ROI following an established cross-validated singular value decomposition (SVD) procedure used in fMRI [45]. For each fold of the data, we decomposed the average (stimuli × voxels) activity matrix from a subset of runs into SVD components, reconstructed the data with varying numbers of components, and selected the dimensionality that maximised the correlation with an independent validation run. These best-fitting dimensionality estimates were averaged across folds. This analysis was performed separately for each rule axis (training stimuli on x- and y-axis). Permutation tests showed that reconstruction correlations were significantly above zero in all ROIs for both axes (**Table S1a**), implying that dimensionality estimates were meaningful.

Estimated dimensionalities for the x- and y-axis did not differ (**Fig. 2a-d** and **Table S1b**) and were themselves significantly correlated in all of regions except for vmPFC (**Table S1c**). Critically, however, we see very different profiles across regions. V1 and PPC showed a modal dimensionality of 1, suggesting that each rule was represented along a neural ‘line’. In hippocampus and vmPFC however, permutation tests showed that the dimensionality was considerably higher across generalizers (HPC>V1: mean=0.695, P<0.001; vmPFC>V1: mean=0.524, P<0.001; HPC>PPC: mean=0.710, P<0.001; vmPFC>PPC: mean= 0.540, P<0.001). Meanwhile, hippocampus and vmPFC did not differ (HPC>vmPFC: mean= 0.171, P=0.1312), nor did V1 and PPC (V1>PPC: mean=0.015, P=0.7157). We also tested whether the projection of stimuli onto the first neural dimension correlated with the order that defined their ground truth treasure locations. We found that it did for V1 and PPC, but surprisingly also for HPC and vmPFC, suggesting that even where multiple dimensions encode the stimuli, the first neural axis carries useful information about the x- or y-location (see **Table S2**, P<0.001 in all four regions and for both x-axis and y-axis).

High- and low-dimensional representations have complementary costs and benefits. High-dimensional representations in which shapes and colours (e.g. red vs. blue, crab vs. duck) are well separated confer flexibility, enabling faster learning in the event that the rule was changed [19]. To verify that higher-dimensional representations facilitated feature individuation, we computed the partial correlation between the estimated dimensionality and the mean decoding accuracy of a linear classifier (**Fig. 2e-f**) taking into account the effect of noise. We found that the two were closely related in both hippocampus (x-axis ρ=0.407, P=0.001 y-axis ρ=0.412, P<0.001) and vmPFC (x-axis ρ=0.592, P<0.001; y-axis ρ=0.314, P=0.0097). Thus, high-dimensional neural codes in hippocampus and vmPFC may support the individuation of specific features.

### Geometry of neural codes during composition

Having established that the representations of individual rules were low-dimensional in V1 and PPC and high-dimensional in hippocampus and vmPFC, next we asked how the representations of rules were geometrically related (**Fig. 3a**). An obvious representational scheme would orthogonalize neural codes for the two features, as their corresponding ground truth outcomes lie at right angles to each other (orthogonal model; **Fig. 3b**). Alternatively, each rule can be viewed as describing the parallel progression along a single mental ‘line’ (parallel model; **Fig. 3c**). Note that in this parallel model there are two possible parallel alignment schemes, one in which the progression from left-to-right on x-axis corresponds to that from top-to-bottom on the y-axis (we call this a ‘top-left’ scheme; **Fig. 3c left**). Conversely, right-to-left on x-axis may map to top-to-bottom progression on the y-axis (a ‘top-right’ scheme; **Fig. 3c right**).

To evaluate whether neural representations were better explained by an orthogonal or parallel model, we conducted cross-validated representational similarity analysis (RSA) using average cosine distances between the vectors for x-axis and y-axis (also known as a parallelism score or PS [5,20], ranging from -1 to 1). PS deviating from 0 in either direction (positive or negative) indicates increasingly parallel axes. We coded the axis locations in a way such that a top-left alignment would have a positive PS (**Fig. 3c left**, corresponding model RDM shown in **Fig. 3d third panel**) and a top-right alignment would have a negative PS (**Fig. 3c right**, corresponding model RDM shown in **Fig. 3d fourth panel**). Next, using a k-fold approach, we regressed the neural RDM of the held-out data against the parallel model RDM and the orthogonal model RDM (**Fig. 3d**). An axis separation model RDM was also included to account for the feature difference between training stimuli from different axes (e.g. training stimuli on the x-axis had the same colour which was different from those on the y-axis) (**Fig. 3d left panel**). We performed permutation tests on the regression coefficients averaged across folds for each model.

In V1, the neural representation was significantly correlated with the orthogonal model (**Fig. 3e**; mean=0.065, P=0.0056). In fact, the whole stimulus space was represented by V1 with two orthogonal low-dimensional features. When we used multidimensional scaling (MDS) to visualise the neural representation of all the stimuli in visual cortex, we saw that it faithfully encoded the correct treasure location, forming a clear 2D grid in neural state space (**Fig. 3i**, see **TableS3** and **Figure S3a** for whole-brain searchlight RSA on 2D representation of all stimuli).

In hippocampus, however, the neural representation correlated with the parallel model (**Fig. 3g**; mean=0.047, P=0.0040). This means that neural codes in the hippocampus represented the x- and y-axes in a high-dimensional but parallel format, with neural responses for the ranked features on the x-axis and y-axis offset by a constant vector in neural state space. Generalizers showed different axis alignment schemes, with some mapping ‘top’ to ‘left’ and ‘bottom’ to ‘right’, some showing the opposite pattern, and the others showing orthogonal axes. To visualize this, we categorized them into different alignment schemes according to the relationship between their PS and the null distribution (**Fig. 3j**) and used MDS to visualize neural geometry for training stimuli in each group (**Fig. 3k**).

This coding scheme is what would be expected if the hippocampus encodes the x- and y-axis with an abstract ‘place’ code, in which the neural population (here, multi-voxel patterns) codes for the progression along the x- and y-axes as if they were linear tracks oriented in parallel (**Fig. 4a**). Because the orientations of the linear tracks can be arbitrary, the mapping may be inverted (giving rise to either a top-left or top-right alignment schemes described above). By contrast, if the neural assembly remapped randomly, then the resulting representations of the linear tracks would be orthogonalized. To illustrate this, we simulated three populations of place cells (100 in each population), each coding for a position on the progression axis (top to bottom or left to right). In different simulations, we controlled for the percentage of cells that remapped across the two linear tracks, with the remaining non-remapped subpopulation had their place fields inverted or kept constant (**Fig. 4b-d**). We show that this model recreates the pattern of parallel, high-dimensional neural coding shown in **Fig. 3k**.

**Figure 4.**
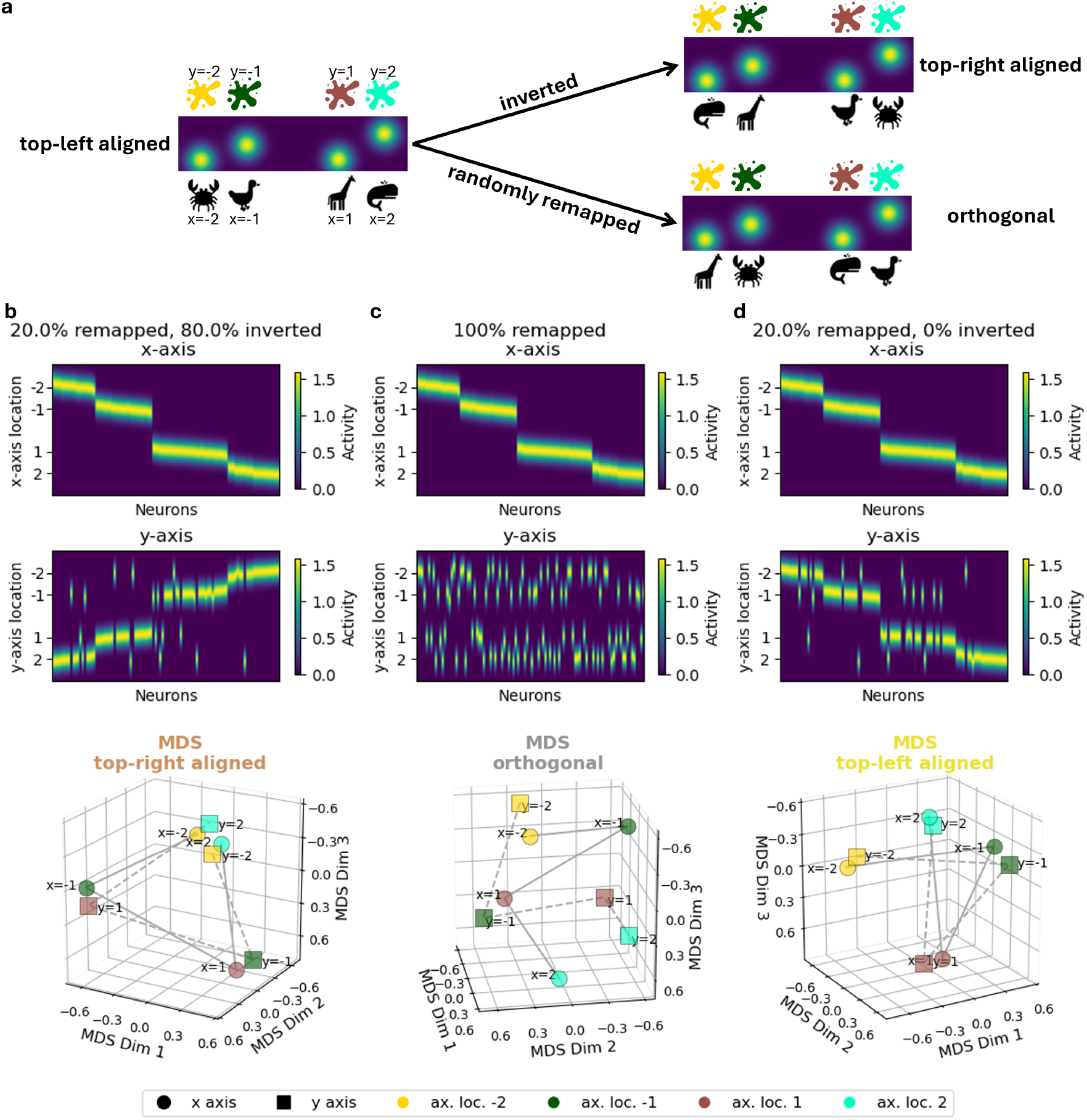
An illustrative model of orthogonal and parallel representation induced by hippocampus remapping. (a) Schematic illustration of the ‘place’ code of four neurons that represent the x-axis and y-axis rules if they were parallel linear tracks. The top shows example neurons responsive at the features associated with the four locations on the y-axis. If there is no remapping, each neuron encoded the features associated with the same location on the linear tracks, learning to a top-left aligned parallel representation (right panel). Note that the axes could be orientated arbitrarily, so the mapping could be inverted, leading to a top-right aligned parallel representation (left panel). If there is random remapping, the population code for the two linear tracks would be orthogonalized (middle panel). (b-d) Three populations of artificial place cells encoding the x-axis and y-axis as two linear tracks with different fractions of neurons that remapped randomly: (b) 20% of the population remapped, the remaining 80% showed inverted mapping across axis; (c) the entire population remapped; (d) 20% of the population remapped, the remaining 80% kept constant. Activity of neurons on each linear track are shown in the top panels. The resulting representations are visualized with MDS in the bottom. Note how the MDS in (b), (c) and (d) resembled the left, middle, and right panels of **Fig. 3k**.

### Neural mechanisms of compositional generalization

Thus far, we have established the representations of individual rules and their geometric relationship to each other. Next, we asked how the representations of rules are combined to enable generalization. A plausible neural strategy is to combine the relevant features in each rule with an additive code, in which the neural patterns for relevant features are combined via vector addition. For example, even if the jade crab has not been seen before, its location can be inferred by adding neural patterns for ‘jade’ and ‘crab’. Previous studies have shown that the neural activity evoked by a compound stimulus resembles the superposition of activity patterns of its individual components [29,46], and does so more strongly than patterns formed by unrelated components [7]. However, a direct test of this class of neural computation is still lacking. To address this, we predicted neural patterns on test stimuli as linear combinations of those obtained on training stimuli. For each test stimulus (e.g. jade crab), we modelled its multi-voxel activity pattern as a weighted linear combination of patterns from “relevant” training stimuli with which it shared a feature (jade elephant and blue crab) and “irrelevant” training stimuli with which it shared none (2 relevant and 6 irrelevant items; the central one was discarded as it is not relevant for any of the test stimuli; see **Fig. 5a**). This regression yielded 8 coefficients (termed “retrieval weights” hereafter) for each test stimulus, forming a 16×8 retrieval pattern matrix (see **Fig. 5b** for two models of retrieval pattern matrix). We used two-fold cross-validation, fitting the retrieval pattern matrix in odd runs and evaluating the fit in even runs, and vice versa.

**Figure 5.**
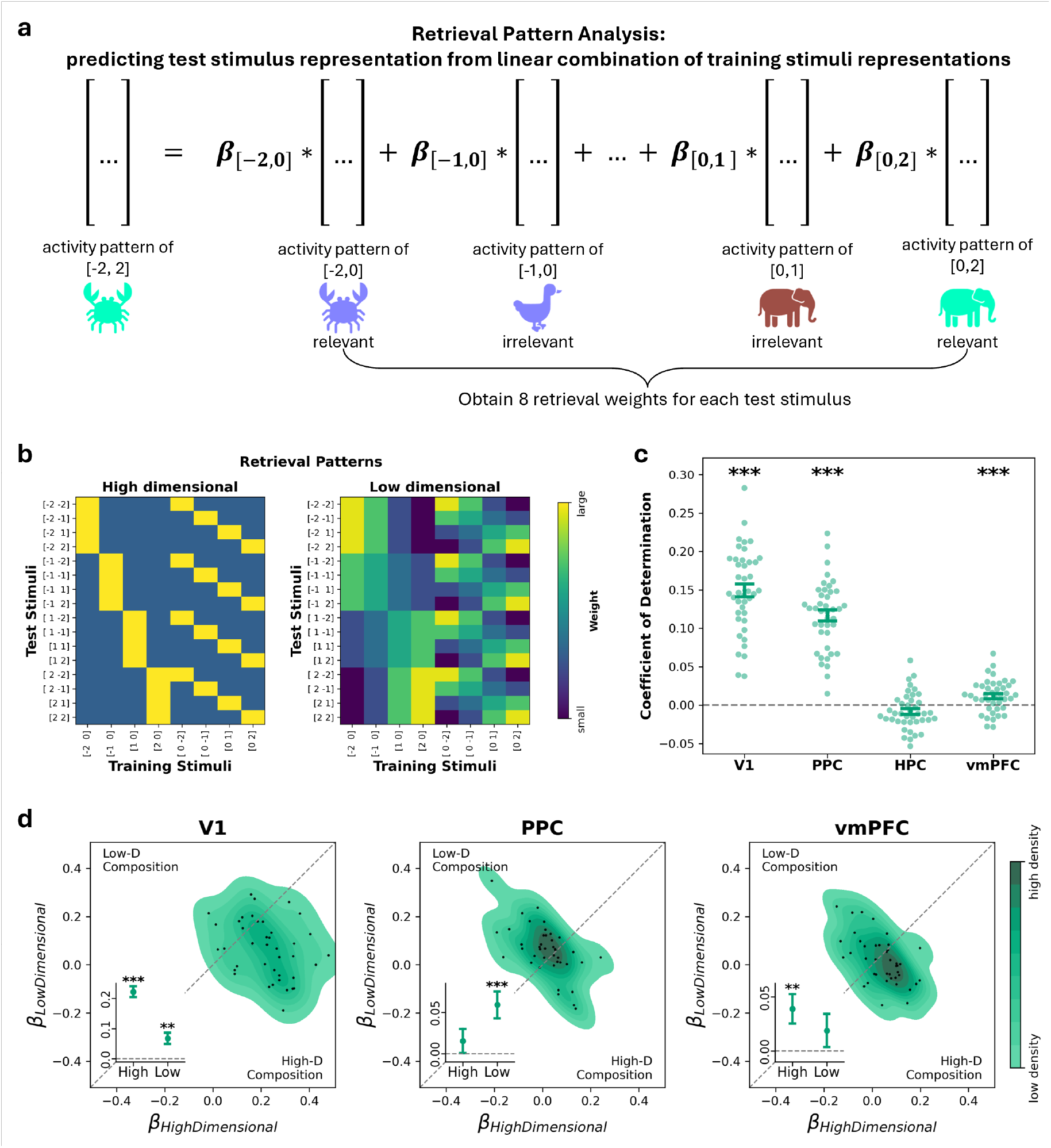
Testing vector addition analysis with retrieval pattern analysis in different ROIs. (a) Schematic illustration of the retrieval pattern analysis. The test stimulus activity pattern was predicted from a (weighted) linear combination of training stimuli activity patterns or pirate positions activity patterns. In this example, the relevant training stimuli/pirate positions at [-2,0] and [0,2] should contribute more to the prediction of test stimulus as [-2,2] compared to the other irrelevant training stimuli. (b) High- and low-dimensional models for retrieval patterns. Top: High-dimensional model indicates that only training stimuli with the relevant features will be used for composition, i.e., a perceptual matching model. Bottom: Low-dimensional model indicates that the irrelevant training stimuli should be weighted by their proximity to the relevant training stimuli, i.e. a distance modulated model. (c) Goodness of fit of the retrieval pattern weights applied to held-out data. Goodness of fit is measured by coefficient of determination. Dots represent individual generalizers. Error bars indicate mean ± standard error. Asterisks indicate results of permutation test against zero: ^***^ P<0.001, ^**^ P<0.01, ^*^ P<0.05. (d) Joint 2D kernel density estimation of empirical correspondence between each retrieval patterns and the high-vs. low-dimensional model, where the colour fill represents the smoothed density using Gaussian kernels. Correspondence is defined as the regression coefficient of a model pattern. Insets show the mean ± standard error of the regression coefficients of model patterns in different ROIs.

If the retrieval pattern matrix yielded above-chance prediction when applied to the held-out data, it meant that for a given region, representations of test stimuli can be reconstructed from those of the training stimuli. We quantified this using the average coefficient of determination across the two folds (note that this statistic is centred on zero under the null, due to the cross validation; **Fig. 5c**). Permutation tests revealed that it was significantly positive in V1 (mean=0.150, P<0.001), PPC (mean=0.117, P<0.001) and vmPFC (mean=0.012, P<0.001) but not in hippocampus (mean = -0.008, P =0.9827). Moreover, vector addition predicts that the relevant training stimuli should contribute more to the prediction than the irrelevant ones. Therefore, we compared their average retrieval weights separately for training stimuli on x-axis (blue animals) and y-axis (elephants of various colours). In the three regions (V1, PPC and vmPFC) with significantly positive R^2^, permutation tests revelated that this was true for both x-axis (**Fig. S4**; V1: mean difference=0.031,P<0.001; PPC: mean difference=0.006, P=0.0387; vmPFC: mean difference=0.006, P=0.0171) and y-axis (V1: mean difference=0.038,P<0.001; PPC: mean difference=0.007, P=0.0031; vmPFC: mean difference=0.006,P=0.0045). Thus, in each of our ROIs except for hippocampus, we saw evidence for a vector addition solution to compositional generalization.

Finally, we tested if the vector addition in V1, PPC and vmPFC occurred in a high- or low-dimensional format. In a high-dimensional code, all features are equidistant from each other, predicting uniformly low retrieval weights for the irrelevant items. We modelled this as a retrieval pattern matrix with entries of 1 for relevant items and 0 elsewhere (**Fig. 5b**, left panel). In a low-dimensional code, however, irrelevant weights should vary with feature proximity, for instance, if brown is spatially closer to jade than green, a brown elephant would have a higher retrieval weight than a green elephant. We derived this low-dimensional model from regression weights obtained by predicting test-stimulus locations from training-stimulus locations—essentially applying the retrieval analysis to ground-truth locations (**Fig. 5b**, right panel). We then regressed the empirical retrieval pattern matrices against the two model matrices (**Fig. 5d**). V1 was significantly correlated with both the high-dimensional and low-dimensional models (high: mean=0.222, P<0.001; low: mean=0.067, P=0.0012). Consistent with our dimensionality analysis, PPC only showed significant correlation with the low-dimensional model (high: mean=0.015, P=0.3144; low: mean=0.059, P<0.001), while vmPFC only showed significant correlation with the high-dimensional model (high: mean=0.039, P=0.0082; low: mean=0.019, P=0.2230). Thus, in V1, PPC and vmPFC composition occurs via vector addition, but in different formats – high-dimensional in vmPFC and low-dimensional in V1 and PPC. Consequently, V1 and PPC showed strong evidence of representing the composites (i.e., the test stimuli) as their corresponding treasure locations (**Fig. S3b, Table S4**). Despite both using low-dimensional formats, V1 and PPC differed in a keyway: V1 encoded all stimuli within a shared low-dimensional reference frame (**Fig. S3a, Table S3a**), whereas PPC maintained a strong separation between training and test stimuli (**Fig. S3a, Table S3b**).

### Space as a scaffold for compositional generalization

In the treasure hunt task, generalizers were required to combine nonspatial features (colour and shape) from two separate training trials to infer the correct treasure location on each test trial. Nevertheless, we wondered whether generalizers might use neural codes for space as a scaffold for making compositional inferences about novel stimuli. In anticipation of this question, after the four runs of the treasure hunt task, we presented participants with a single run of an additional ‘spatial localiser’ task (**Fig. 1d**). In the localiser task, a small pirate icon appeared at one of the training stimulus locations in the circular arena on each trial. Participants performed a cover task where they were asked to press a button when the current pirate position was the same as the previous trial.

First, we investigated whether the four ROIs encoded the spatial position of the pirate. To do so, we performed RSA analyses in each ROI by computing the spearman’s rank correlation between the neural representation and a spatial representation model (**Fig. 6a**). Permutation tests showed that all four ROIs showed significant correlation with the spatial model (**Fig. 6b**; V1: ρ=0.745, P=0.0002 ; PPC : ρ=0.1162, P=0.0016; HPC: ρ=0.088, P=0.0110; vmPFC: ρ=0.077, P=0.0252). To further validate this, we repeated the same analysis in a whole-brain searchlight RSA. Only V1 and PPC survived cluster correction in the whole-brain searchlight (**Table S5** and **Fig. S3c**). Therefore, the spatial representation of pirate positions was strongest in V1 and PPC, and the effect was weaker in HPC and vmPFC. MDS visualization of the representation in V1 using MDS faithfully reconstructed the pirate positions in the localiser task (**Fig. 6c**). Next, we asked if this spatial code is shared with the low-dimensional code in V1 and PPC during the treasure hunt task. An intuitive way to quantitively test this is to repeat the previous retrieval pattern analysis. However, the procedure required fitting retrieval weights using data from the same run, which is not applicable here as the data for treasure hunt task and localiser task was collected in different runs. Therefore, we performed a cross-task representation similarity analysis (cross-task RSA) instead.

**Figure 6.**
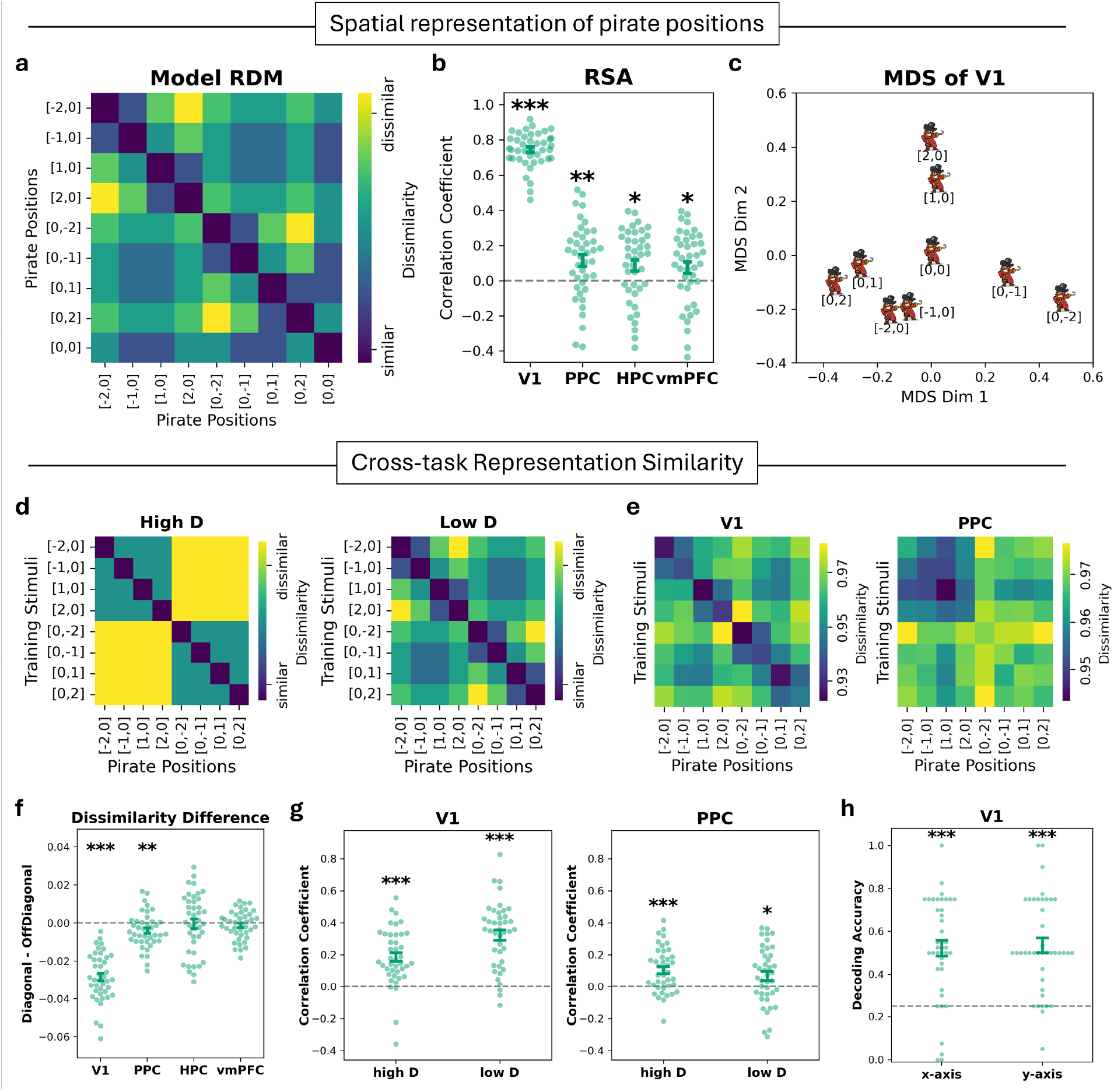
Low-dimensional representation in V1 and PPC is shared with spatial representation. (a) Model RDM for spatial representation of the pirate positions. (b) Correlation with the spatial model in the four ROIs. (c) MDS visualization of V1 representation of the pirate positions. (d) High and low dimensional model RDMs for cross-task RSA. Each entry reflects the neural distance between a stimulus and a pirate position. (e) The symmetrized empirical cross-task neural RDM observed in V1 and PPC. (f) Difference in neural dissimilarities between pairs where pirate position matched training stimulus location (diagonal entries of cross-task RDM) and pairs where pirate position did not match training stimulus location (off-diagonal entries). (h) Cross-task decoding accuracy. We trained logistic classifiers in the treasure hunt task to predict the test stimulus location on a given axis and evaluated the classifier using data from the localizer task. Translucent dots represent the decoder’s mean evaluation accuracy across 20 random seeds for each participant. Error bars represent mean ± standard error. Dashed grey lines represent chance level.

**Figure 7.**
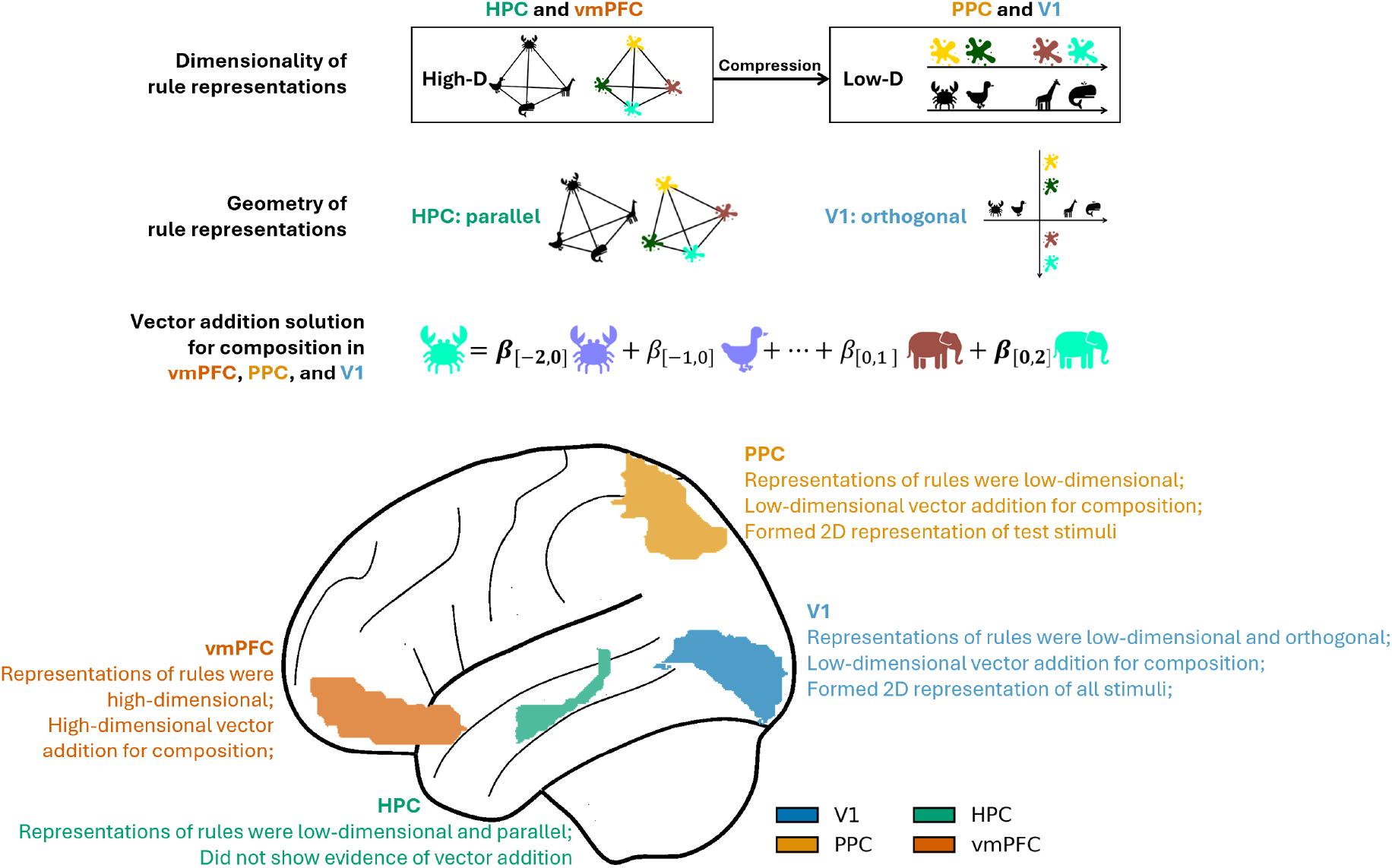
Summary of results. We studied the neural representations of rules and the composition of rules that support successful inference on novel combinations (i.e., test stimuli). Across four ROIs, we saw variations in the dimensionality and geometry of the rule representations: representations of individual rules were high-dimensional in HPC and vmPFC and low-dimensional in PPC and V1. In HPC, representations of the two rules were parallel in a way that reflected the abstract relational structure shared between rules. Meanwhile, vmPFC, PPC, and V1 showed evidence of a vector addition solution for compositional generalization.

For each ROI, we constructed a cross-task neural RDM with entries reflecting the neural distance between a training stimulus and a pirate position. Since each pirate position corresponds to one of the training stimulus locations, we first tested whether matching stimulus-position pairs (diagonal entries in the neural RDM) were represented more similarly than non-matching pairs (off-diagonal entries) (**Fig. 6f**). This was true in V1 (mean difference=-0.028, P=0.0002) and PPC (mean difference=-0.004, P=0.0076), but not HPC (mean difference=-0. 0004, P=0. 8787) and vmPFC (mean difference=-0. 0012, P=0.3332). Next, we conducted a follow-up analysis to examine whether the dissimilarity of non-matching pairs was better explained by a high-dimensional or low-dimensional model. Similar to retrieval pattern analysis, high-dimensional model predicts a difference between matching and non-matching pair, but no difference among non-matching pairs, and low-dimensional model predicts neural distance modulated by spatial proximity. To do so, we first symmetrized the asymmetric cross-task RDMs by averaging across the diagonal (**Fig. 6e**). We then regressed the lower triangle of each symmetric RDM onto the model RDMs shown in **Fig. 6d**. Both V1 and PPC showed significant correlations with both the high-dimensional model (V1: mean=0.187, P=0.0002; PPC: mean=0.106, P=0.0002) as well as the low-dimensional model (V1: mean=0.325, P=0.0002; PPC: mean=0.070, P=0.0162; **Fig. 6g**).

Finally, we ran an ambitious cross-task decoding analysis to validate if classifiers trained to predict locations can generalize across task. We trained a logistic regression classifier using data from the treasure hunt task to decode the ground-truth location on a given axis from the activity patterns of test stimuli. And then, we evaluated whether the decoder can be applied to predict the pirate’s position on the same axis in the localiser task (**Fig. S5a**). We performed this analysis separately for the x-axis and y-axis. While PPC did not show above-chance cross-task decoding performance (x-axis: mean=0.243, P=0.852; y-axis: mean 0.273, P=0.504), V1 showed strikingly strong cross-task decoding accuracy (x-axis: mean=0. 522, P=0.0002; y-axis: mean=0.536, P=0.0002; **Fig. 6h**, see **Fig. S5c** for performance in other ROIs and see **Fig. S5d** for performance of classifiers trained on training stimuli). This suggests that, even though test stimuli were presented at the centre of the screen, V1 encode them as if the generalizers were ‘seeing’ the stimuli in the corresponding ground-truth treasure locations.

## Discussion

We studied the dimensionality and geometry of neural codes, obtained from BOLD multivoxel pattern activity, in participants who successfully performed a compositional generalization task (‘generalizers’). We observed some surprising differences between brain regions. In hippocampus and vmPFC, neural codes were high-dimensional and allowed for feature individuation. In vmPFC, PPC and V1, neural activity patterns on test trials could be predicted as a linear combination of training trials. In vmPFC this combination occurred in the space of individual features, whereas in PPC and V1 it was grounded in the two-dimensional response space and compositional neural codes overlapped with representations of space, recorded from the spatial localiser task.

Our results are consistent with the stimuli (coloured shapes) initially being represented in the hippocampus in a high-dimensional format that maximises feature individuation [47], i.e. the capacity to flexibly distinguish red items from both blue and green via a linear readout, irrespective of their proximity in outcome space. This would be useful if the system wished to re-combine the representations (shapes and colours) in a different way, for example if we subsequently changed the mapping between colours (or shapes) and ground truth locations on the x-axis (or y-axis) [19]. In the hippocampus, however, we additionally observed a curious phenomenon: the high-dimensional neural axes for each feature (colour and shape) lay parallel to one another in neural state space. This suggests that the hippocampus factorises the two features but represents them on a common high-dimensional trajectory, rather like an (idealised) population of place cells might represent two parallel linear tracks, without remapping. Such a representational format aligns with the hippocampus’s established role in encoding latent relational structure [27,48] and in forming conjunctive codes that bind items to their roles within a structured representation [7,49]. We note that generalizers heterogeneously associated the x- and y-axis with different mapping rules (e.g., top-left; bottom-right). We are unclear as to the purpose of this coding scheme, but it was prominent and highly reliable in our data.

We see evidence that this flexible, high-dimensional format is conserved in vmPFC, which receives prominent projections from the medial temporal lobe, including hippocampus. However, in vmPFC (unlike in hippocampus) we see that test items (e.g. jade crab) are represented as a linear combination of the (high-dimensional) neural representations of ‘jade’ and ‘crab). This makes this region a candidate for composing elementary features into a novel representation, which is consistent with the finding that composite stimuli (e.g. an avocado and raspberry milkshake) can be neurally primed by individual elements (avocado and raspberry) in this region [30]. However, the vmPFC did not contain information about the spatial location at which the outcome would be found for this novel composite item, and the neural pattern for ‘jade crab’ could not be composed from neural vectors that coded for the x- and y-locations corresponding to ‘jade’ and ‘crab’.

In the PPC, by contrast, we saw that composites were encoded in a low-dimensional format that retained the 2D Euclidean geometry of the outcome space. This means, for example, that if red, green and blue stimuli signalled that outcomes would be found on the left, middle, and right of the response arena, then neural patterns evoked by red and green cues would show greater overlap than those for red and blue cues. This 2D representation was neurally aligned with that recorded during a spatial localiser task, in which potential outcome locations were systematically highlighted. In other words, whilst composition in the vmPFC occurred in the high-dimensional space, in PPC it was reformatted to be maximally useful for the response. This is consistent with the older theory of the neural circuitry underlying decision-making, which argues that vmPFC encodes response-independent contingencies, whereas parietal cortex maps signals into a response-consistent frame of reference [50]. Notably, this transformation into response space was observed only for the test stimuli, not for the training stimuli. Additionally, we found that training and test stimuli were strongly segregated in PPC. The basis of this separation remains unclear, but it may point to a potential role for PPC in situations that require inference.

Finally, by far the strongest low-dimensional code was observed in the primary visual cortex. Unlike PPC, V1 encodes all stimuli in the same low-dimensional space, while also maintaining a strong feature-based representation. We did not anticipate this result. However, we think it is likely that the visual cortex contributes to the maintenance of spatial information about the correct location during the stimulus presentation period, consistent with its known role in working memory [51–53] and anticipating the spatial location of events in the future [54]. It may well be that, given an entire day to practice the task, generalizers learn a relatively automatised strategy to direct their gaze to the corresponding locations that allows them to encode and maintain the response location. However, we cannot rule out the possibility that the visual cortex contributes actively to the compositional process itself.

## Methods

### Participants

In total, we collected two cohorts of participants with 30 participants in each cohort. We preregistered the participant classification criteria, exclusion criteria and the hypothesis before collecting and analysing the data of the second cohort (https://osf.io/9a6r7). Four participants who showed inconsistent behaviour in the scanner were excluded. We classified the remaining 56 participants based on the preregistered criteria. Based on the classification results, we reported all our analysis based on the data from the 41 generalizers who successfully learned and solved the task. Participants were recruited from University of Granada and were paid a base rate of 28 euros with performance-dependent bonus of up to 10 euros. The behavioural experiments were approved by the Medical Sciences Research Ethics Committee of the University of Oxford (R50750/RE001) and the neuroimaging experiment was approved by the Research Ethics Committee at the Research Centre for Mind, Brain and Behaviour at the University of Granada.

### Design

In the main treasure hunt task, participants were instructed to find treasure on a virtual island based on cues. Cues were coloured animal shapes. The hidden rule was that each feature of the cue (colour or shape) indicated the treasure location on the x- or y-axis in cartesian coordinates (**Fig. 1a-b**). The mapping from a perceptual feature to the corresponding continuous spatial axis was arbitrary and randomized across participants.

In each trial, participants first saw the cue appear in the centre of the screen for 1 second. After the cue disappeared, a circular arena appeared, and participants had to move the pirate to the treasure location with four keys that corresponded to movement in the cardinal directions. The response stage was self-paced; participants could only proceed to the next screen once they had submitted a response for the current trial. To familiarise participants with the layout of the controller in the scanner, we fixed the mapping from fingers to directions (index-middle-ring-pinky corresponded to left-up-down-right). To allow for error tolerance, we define a reward zone with radius *radius*_*reward*_ around the correct location. Reward zones of different stimuli did not overlap. Responses within this zone earn a reward, with higher rewards for greater precision: those falling within the first, second, and third thirds of the radius receive bonuses of 5, 2, and 1, respectively.

In training blocks, feedback on the correct treasure location was presented for 2s (**Fig. S1b**). In test blocks, no feedback was provided (**Fig. S1c**). During feedback, participants saw their response location, the correct location, as well as the three concentric circles indicating the three levels of possible reward. Importantly, participants were only exposed to the 9 stimuli in training blocks (**Fig. 1a**), and the remaining 16 unseen colour-shape combinations were used as test stimuli in the test blocks (**Fig. 1b**). Therefore, participants had to make zero-shot inferences on these unseen combinations to solve the task.

To make sure we could have as many generalizers as possible, we picked the axis-aligned blocked training curriculum from a previous study [8] (**Fig. S1a**). In this curriculum, the training stimuli locations were aligned to the cardinal axes and constituted a “+” in the centre of the map, i.e., stimuli in the middle row and column. These stimuli were presented in a blocked fashion in a training block. Participants first saw stimuli on one axis, then the stimulus at the centre location where the axes crossed, and then stimuli on the other axis. The order in which the axes were presented alternated across training blocks. In the test blocks, test stimuli (off-axis) were presented randomly.

During scanning, participants were evaluated on all stimuli in the treasure hunt task without feedback. Participants first saw the cue appear in the centre of the screen for 1 second. In one third of the trials in each run, this was immediately followed by a response screen where participants were allowed a maximum of 8 seconds to move the pirate to the treasure location. In the remaining two thirds of the trials, no response was required and the experiment proceeded to the next trial. A jittered inter-trial interval (ITI) was used (mean = 3 s, range = 1–5 s). All stimuli were presented randomly.

To obtain the neural representation of spatial locations, we designed another localizer task for scanning. In each trial, participants saw the pirate appear at a location within the arena and had to indicate whether the pirate position was the same as on the previous trial using a keypress. The pirate position could be any of the nine training locations.

### Procedure

The study consisted of two sessions (usually two days, with 9 out of 60 participants completing both sessions on the same day). Participants completed a pretraining session online before coming to the refresher and scanning session in person (**Fig. S1a**). In the pretraining session, participants completed 6 train-test cycles. Each cycle had two training blocks followed by a test block. In the scanning session, participants completed a refresher of three training blocks before going into the scanner. In the scanner, participants completed four runs of the treasure hunt task followed by one run of a localizer task. Each run of the treasure hunt task had 3 repetitions per stimulus (25×3=75 trials per run). Due to time constraints, participants were only required to move the pirate in one of the repetitions per stimulus. No feedback was provided, and all trials were randomly presented. The localizer task had 20 repetitions per training location (9×20 = 180 trials). Our analyses focused on the cue presentation stage in both tasks.

### fMRI data collection

We acquired T2-weighted functional images on a 3T SIEMENS MAGNETOM Prisma scanner with a 64-channel head coil and a multi-band acceleration factor of 2. We acquired 50 slices, 2.5mm thick with: repetition time (TR) = 1730ms, echo time (TE) = 30ms, flip angle = 66°, field of view (FoV) = 210 mm, voxel size = 2.5×2.5×2.5 mm^3^. To correct for spatial distortion, a field map was acquired with dual echo-time images covering the whole brain with the following parameters: TR = 520ms, TE1= 4.92ms, TE2 = 7.38ms, flip angle = 60°, FoV = 210mm, voxel size = 3.0×3.0×2.5 mm^3^. A T1-weigted structural image was acquired with the following parameters: TR = 2250ms, TE= 4.18ms, flip angle = 7°, FoV = 256mm, voxel size = 1×1×1 mm^3^.

### Participant classification and exclusion

In the main text, we reported the analysis of we reported all our analysis based on the data from the 41 generalizers who successfully learned and solved the task. The classification was based on the previous work [8], we defined participants who could correctly position the training stimuli as learners and those who failed as non-learners. We further defined participants who could correctly position the test stimuli as generalizers and those who failed as non-generalizers. We classified participants based on their performance in the scanner using a metric called log-likelihood ratio (LLR). It operates under the assumption that that the response of a generalizer would follow a binormal distribution around the true treasure location (ground truth model). This was compared against a null model where the participant responded in any location on the island equally likely. LLR was computed to describe whether the probability that a participant’s response followed the ground-truth model (*P*_*binormal*_) exceeded the probability that they were responding randomly (*P*_*uniform*_).

To calculate these response probabilities, the arena was discretised into grids, and the normalized probability of response in each grid was obtained for each model. First, we defined the discretised probability mass function as the probability of response falling into each grid. For each grid in the circular arena (*x*^2^ + *y*^2^ < 2), we calculated the unnormalized probability (i.e., does not sum up to one) for each model as follows.

The uniform model was:

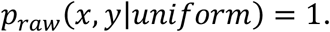

The binormal model was:

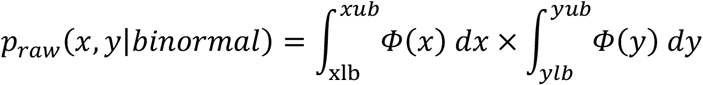

where *xlb, xub, ylb, yub were* the lower and upper bounds of the grid *x, y* falls in. *Φ* was the probability density function of the normal distribution 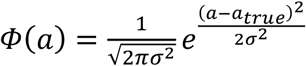, where was *a*_*true*_ the correct x/y coordinate, and we set 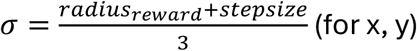, where *radius*_*reward*_ was the radius of the circular reward zone, and *s*_*tepsize*_ represented the precision of movement (change in position induced by one key stroke).

Next, we normalized the probabilities for each grid so that they sum up to 1:

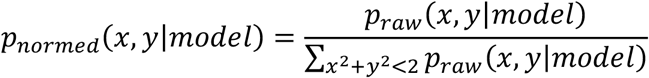

Finally, a lapse rate (*λ*) of 0.01 was added to allow for a small degree of randomness in the response:

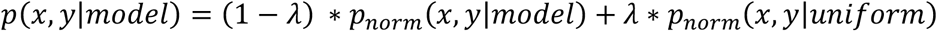

Now that we obtained the normalized probability of response in each grid for each model, the LLR of any given trial *t* could be calculated:

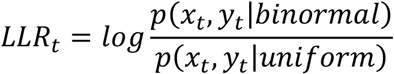

Over *N* trials, the average LLR was computed as:

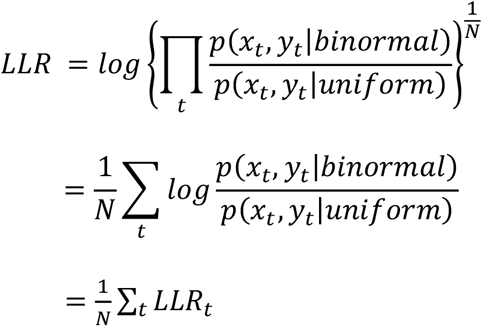

*P*_*binormal*_ > *P*_*uniform*_ would lead to *LLR* > 0. If the *LLR* of training stimuli was greater than zero (*LLR*_*train*_ > 0), the participant would be classified as a learner. If the *LLR* of test stimuli was greater than zero (*LLR*_*test*_ > 0), the participant would be classified as a generalizer. Importantly, we computed the *LLR*_*train*_ and *LLR*_*test*_ for each participant during their last block of pretraining and during scanning. Because there were four runs of scanning, we first computed the average response to each stimulus across all four runs for each participant, then computed *LLR*_*train*_ and *LLR*_*test*_ on the averaged response map for each participant. We classified participants based on their performance in the scanner. We further excluded 4 participants who were learners or generalizers in the last block of pretraining but behaved like non-learners or non-generalizers in the scanner (*LLR*_*train*_ > 0 in pretraining, *LLR*_*train*_ < 0 in scanner; or *LLR*_*test*_ > 0 in pretraining, *LLR*_*test*_ < 0 in scanner, see Figure S3-2). This procedure yielded 41 participants classified as learner-generalizer (referred to generalizers hereafter) across both cohorts and they were all included in the reported analyses.

### Preprocessing of fMRI data

All fMRI data was pre-processed in SPM12. For each participant, we acquired data from 5 functional runs (EPI images), a fieldmap and a high resolution T1 image. First, we performed auto-reorientation so that all the acquired images were roughly in the same space as the MNI152NLIN6Asym template [55,56]. Second, we calculated the voxel displacement map (VDM) from the fieldmap image. We then corrected signal distortion based on the calculated VDM image and realigned the images to the first volume of the first run. Third, we coregistered the T1 image to the mean EPI image of the first functional run. Fourth, we performed segmentation and normalization on the coregistered T1 image using the default tissue probability maps in SPM. Finally, the normalization parameter estimated using the coregistered T1 image were used to normalize all the EPI images. Normalised functional images were then used to perform the multivariate analysis without smoothing.

We used the estimations of the six head motion parameters (three translation parameters on x,y,z planes and three rotation parameters: pitch, roll and yaw) generated by SPM during realignment to calculate framewise displacement [57]. First, we converted rotational parameters from radians to millimetres by calculating displacement on the surface of a sphere of radius 50 mm. Then, we calculated the derivatives for the six parameters (e.g. for framewise head movement on x: 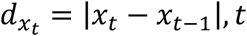, *t* being the current timepoint). Finally, for each TR, we defined framewise displacement as the sum of the six derivatives. If participants demonstrated excessive head motion in the scanner (more than 10% framewise displacement greater than 1 voxel size), they would be excluded from further analysis. No participants were excluded due to excessive head movement.

### Preparing Data for Multivariate Analyses

To understand the neural geometry that supports compositional generalization, we adopted multivariate approaches to analyse activity patterns induced by stimuli in the tasks. All multivariate analyses were conducted in Python, except for the group-level analyses for searchlight results.

#### Anatomical ROIs

We anatomically defined our regions of interest (ROIs), ventromedial prefrontal cortex (vmPFC) and hippocampus (HPC) based on previous literature suggesting their role in inference. We further included early visual cortex (V1) as a control region. The vmPFC and HPC masks were generated from the AAL parcellation [58]. The V1 mask was generated from the HCP-MMP1.0 parcellation [59] that was projected onto the MNI152NLIN6Asym template [60]. Before running the multivariate analyses (in ROI or in a searchlight sphere), we obtained whitened activity patterns for each stimulus as described below.

#### Obtaining activity patterns

To obtain the activity pattern of each stimulus, we ran a first-level GLM to estimate the contribution of each stimulus to each voxel’s activation level following the LSA approach [61]. The onset events of each stimulus in each run were modelled as a separate regressor in this GLM. We ran this GLM for each participant on their preprocessed fMRI data in SPM12. We extracted the estimated coefficients for each stimulus as the raw (unwhitened) activity pattern matrix ***B*** where each row *b*_*k*_ is the unwhitened activity pattern of a stimulus in one run. Then we performed multivariate noise normalization (‘*whitening’*) on the raw activity pattern [62,63]:

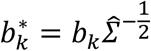

where the voxel-by-voxel covariance matrix 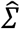 is estimated from the residuals of the GLM using Oracle Approximating Shrinkage Estimator (OAS) implemented in the scikit-learn package [64]. Because the covariance matrix 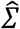 is Hermitian, so we can obtain the whitening matrix 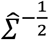 via the eigen decomposition of 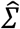.

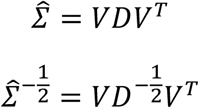

The whitened ***B***^*^ were used for the following multivariate analyses.

### Cross-validated dimensionality estimation of the representations of individual rules

To quantify the dimensionality of the representation of each rule, we examined the dimensionality of non-centre training stimuli representation on each axis separately with four-fold cross-validated singular value decomposition (SVD) [45].

Let 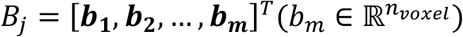 denote the activity pattern matrix of *m* stimuli from a run *j* that serve as the held-out subset. The remaining three runs were iteratively partitioned into a fitting subset (*B*_*fit*_, two runs) and a validation subset (*B*_*val*_, one run).

For each partition, the mean activity pattern of the fitting set, 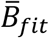, was computed and decomposed via SVD:

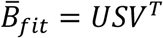

where *U* = [***u*_1_, *u*_2_, *u*_3_, *u*_4_**] and 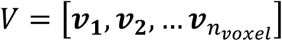 were the left and right singular vectors, and the diagonal entries of *S*, denoted *s*_*i*_, are the singular values. We reconstructed 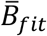 at different dimensionalities *k* ∈ {1,2, …, *m*}. The reconstruction at dimensionality *k*, denoted 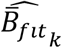, was given by:

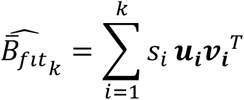

Then, we computed the Pearson correlation *r*_*p*_(*k*) between the reconstructed data and the validation data for the current split.

Next, we selected the dimensionality 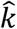 that maximized the mean correlation across all *P* possible fitting/validation splits:

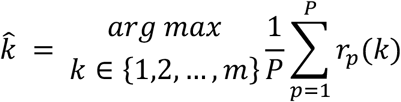

This 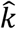 was taken as the estimated functional dimensionality for the held-out run *j*. To obtain a metric for reconstruction quality, we averaged the activity patterns of fitting and validation subset (denoted 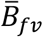), computed its 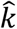 dimensional SVD reconstruction 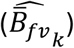, and computed the Pearson correlation between 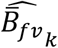 and the held-out run *B*_*j*_. This correlation served as a metric of reconstruction quality and was also used as a proxy for signal quality (i.e., noise) when we computed the partial correlation between the dimensionality estimates and the decodability of stimuli features.

### Geometry analysis on the alignment between representations of individual rules

#### Computing parallelism score (PS)

To run our cross-validated representation similarity analysis, we first estimated axis alignment by computing the parallelism score. Let *X* = [**x**_**−2**_, **x**_**−1**_, **x**_**1**_, **x**_**2**_] *and* 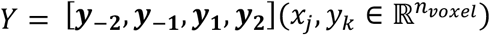 be the activity pattern matrix for all the non-centre training stimuli on the x and y axis respectively for a single participant. Within each axis, we computed the coding direction between a pair of locations. For instance, the coding direction between location ***i, j*** on the x axis was ***d*x**_***ij***_ = **x**_***i***_ **− x**_***i***_, and the same for y: ***dy***_***ij***_ = ***y***_***i***_ **− *y***_***i***_. Then we computed the cosine similarity between ***d*x**_***ij***_ and ***dy***_***ij***_ to measure the alignment of the coding directions between ***i, j*** on the x and the y axis. Finally, we obtained the parallelism score as the average cosine similarity across all possible location pairs.

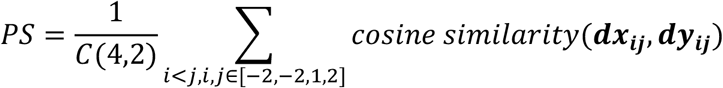

Perfectly orthogonal axis representations would lead to a PS of 0. A negative PS indicated that the top of the y-axis was aligned to the right of x-axis, whereas a positive PS indicated top-left alignment between y-axis and x-axis.

#### Four-fold cross-validated RSA

In this analysis, the training stimulus activity patterns from two runs (fitting set) were averaged to calculate the PS. Then, we ran an RSA with the averaged activity pattern across the other held-out runs (evaluation set) where we regressed the neural RDM of the training stimuli on three model RDMs (**Fig. 3d**): axis separation, parallel and orthogonal. Axis separation indicated that training stimuli on the same axes were represented more similarly to each other than to those from different axes. The orthogonal RDM was generated from the distance between the ground truth location on the map. The parallel RDM was generated based on the PS from the fitting set: PS>0 indicated a top-left alignment (**Fig. 3c** left panel and **Fig. 3d** third panel) such that the top of y-axis (y=-2) was more similar to the left of x-axis (x=-2), and that y=-1 was more similar to x=-1, and so on. PS<0 indicated a top-right alignment (**Fig. 3c** right panel and **Fig. 3d** fourth panel) such that the top of y-axis (y=-2) was more similar to the right of x-axis (x=2), and that y=-1 was more similar to x=1, and so on. We repeated this process across all possible fit-evaluation splits and computed average regression coefficients across these splits as the metric of similarity between neural representations and the parallel or orthogonal model RDM.

#### Classification into different alignment schemes for visualization

To visualize the geometry of rule representations in HPC, we classified generalizers into different axis alignment schemes by comparing their PS to a null distribution. We generated a null distribution of PS at the group level with the following procedure. We took the activity pattern matrix of all the non-centre training stimuli (each column was the activity pattern of one training stimulus, columns ordered by axes and the locations on each axis: from x=-2 to x=2, then from y=-2 to y=2). For each participant, we randomly shuffled the labels of the columns (i.e., randomly assigned the activity patterns to stimuli), then computed the PS for this shuffled matrix and this participant. And then, we averaged across all generalizers to obtain an average PS for one shuffle. We repeated this process 10,000 times and obtained a group null for PS, denoted as *PS*_*HPCnull*_.

Generalizers’ axis alignment schemes were classified by comparing their PS against the group null in the hippocampus ROI. Let *PS*_*HPC,j*_ be the PS for participant j:

1. If *PS*_*HPC,j*_ was significantly greater than the null 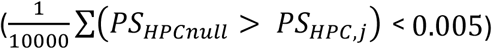, then the participant was classified as *top-left aligned*.
2. If *PS*_*HPC,j*_ was significantly smaller than the null 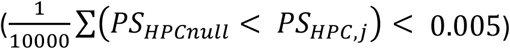, then the participant was classified as *top-right aligned*.
3. Otherwise, the participant was classified as *orthogonal*.

#### Illustrative model

To illustrate how place-like code could give rise to the observed axis alignment, we simulated three populations of place cells, each with 100 neurons. Each neuron had a preferred location on the x-axis (-2, -1, 1 or 2) and a preferred location on the y-axis (-2, - 1, 1 or 2). In each population, a percentage of neurons may remap. We randomly sampled the preferred location on the x-axis and that on the y-axis for the remapped subset of neurons. The remaining non-remapped subset either has the same preferred location on both axes or has inverted locations (e.g. -2 for x-axis and 2 for y-axis). In the first population (**Fig. 4d**), 20% of the population remapped, the remaining 80% kept constant 。 In the second population, 20% of the population remapped, the remaining 80% showed inverted mapping across axis (**Fig. 4b**). In a third population (**Fig. 4c**), the entire population remapped.

### Retrieval pattern analysis

To test the vector addition hypothesis, we ran regressions to predict the activity pattern of test stimuli as linear combinations of activity patterns of training stimuli (**Fig. 5a**). To avoid overfitting, we averaged the activity patterns of a given stimulus across odd runs and across even runs to obtain an odd-even split for cross-validation.

Regression weights fitted with the averaged patterns from the odd split were applied to the even split to validate the regression model fit and vice versa. Let *X*_*tr,odd*_, *X*_*tr,even*_ be the activity pattern matrix of training stimuli for odd and even splits respectively, and *X*_*te,odd*_, *X*_*te,even*_ denotes the activity pattern matrix of test stimuli for odd and even splits respectively. Each row of an activity pattern matrix (denoted by *x*_*te,odd*_, *x*_*te,even*_) is the activity pattern of a stimulus. We obtained two weight matrices *W*_*odd*_ and *W*_*even*_ in which each row (denoted by *w*_*odd*_, *w*_*even*_) is a set of ordinary least square estimates obtained for predicting a test stimulus:

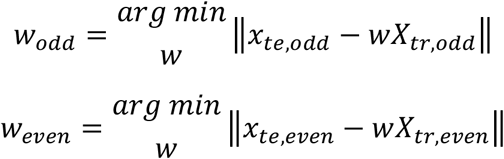

where ‖∙‖ indicated L2-norm of a vector. The final retrieval pattern matrix was computed by averaging *W*_*odd*_ and *W*_*even*_. Intercepts were also fitted.

Model fit was evaluated on the held-out data by applying the weights to make prediction about test stimuli and calculating the coefficient of determination (denoted *R*^2^ hereafter). Note that *R*^2^can be negative because the held-out data were not seen during fitting. If the *R*^2^ was significantly above zero, it indicated that the prediction in the held-out data was successful. Specifically, for odd and even split, we computed the mean *R*^2^ across prediction of all test stimuli :

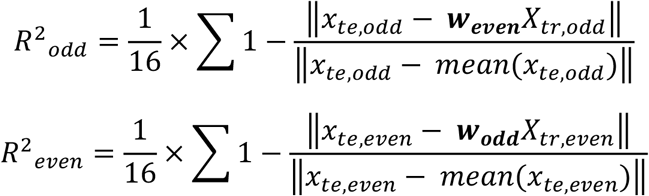

The final *R*^2^ was computed by averaging *R*^2^*_odd_* and *R*^2^*_even_*.

Moreover, the vector addition hypothesis implied that the relevant stimuli contributed to the prediction and more so than the irrelevant stimuli. Therefore, we computed the mean weights for the relevant stimuli and irrelevant stimuli separately and for each participant. Then we tested if the mean weight for relevant stimuli was greater than zero and whether the mean weight for relevant stimuli was greater than the mean weight for irrelevant stimuli at the group level. If the prediction was successful, and both tests were satisfied, it meant that for a given region, composition happened through vector addition.

After these sanity checks, we explored whether vector addition happened in high or low dimensional space. To this end, we first defined the retrieval weight matrix for test stimuli by averaging the weights obtained from odd and even splits. Then, we designed two model retrieval patterns that corresponded to composition in high and low dimensional space and compared the obtained retrieval pattern matrix against the models. For high-dimensional composition, this process happened through perceptual feature matching, i.e., to make an inference about a blue crab, one would only retrieve information from those training stimuli that were either blue or a crab. This resulted in a high-dimensional model retrieval pattern where only the relevant training stimuli would have high weights. If composition happened in low dimensional space, the retrieval weights for irrelevant training stimuli would be further modulated by the distance to the relevant training stimuli whereby closer distance yielded larger weights. Since these two model matrices were correlated, we entered both models in the same regression to predict the obtained retrieval pattern matrix.

### Decoding analysis

All decoding analyses reported were performed using a linear logistic regression decoder with l2 penalty. To make sure the obtained results were not sensitive to random seeds, we repeated all decoding analyses with 20 random seeds and reported the average across random seeds. All decoding analyses were performed with the scikit-learn package [64].

#### Decoding features of training stimuli

This analysis accompanied the dimensionality analysis to address the role of dimensionality in maintaining feature separability. We conducted two separate decoding analyses, one focussing only on the features on the x-axis (e.g. colour) and one on the features on the y-axis (e.g. shape). To avoid overfitting to run-specific noise, we adopted the repeated stratified k-fold scheme for cross-validation. Take x-axis feature decoding as an example, for each participant, activity patterns of non-centre training stimuli on the x-axis from all four runs were randomly divided into four splits (disregarding run identity). The logistic regression decoder was fitted on three out of the four splits and evaluated on the held-out split. Within the fitting data, we further performed hyper-parameter tuning with repeated stratified three-fold cross-validated grid search to look for the best performing penalty parameter. The decoder was refit to all fitting data with the identified penalty parameter and evaluated on the held-out fourth run. This process was repeated four times where each run served as the held-out run once. Decoding accuracy was computed as the average accuracy on the held-out run across the four repeats.

#### Cross-task decoder generalization (from treasure hunt task to localizer task)

To examine whether the representation of stimuli in the treasure hunt task was shared with the location representation, we evaluated the performance of discrete position decoders on the localizer task after fitting the decoder using the treasure hunt task stimulus representations (**Fig. S5a-b**). We performed the analysis for x- and y-axis features separately and for training stimuli and test stimuli separately. Take x-axis location decoding trained on test stimuli as an example, for each participant, we took the activity patterns of the test stimuli and fitted a logistic regression decoder with a hyper-parameter tuning procedure similar to that described in the previous section. After that, we applied the decoder to the localizer task data. Specifically, when the logistic regression is trained to predict x-axis location, we only evaluated it on the four pirate positions on the x-axis, i.e., in the middle row. This was because pirate positions in the middle column occupied an x-axis location that was never seen by the decoder during fitting. For a similar reason, when fitting an x-axis location decoder using the training stimuli, we only used the four stimuli in the middle row as well. The same process was repeated for the y-axis feature.

### Representational Similarity Analysis (RSA)

In representational similarity analyses, we first calculated the representational dissimilarity matrices (RDM) for the neural data. For each searchlight sphere or ROI, we took the whitened activity pattern matrix, and for each pair of stimuli, we calculated the neural dissimilarity using correlation distance (1 minus Pearson correlation). Then, we quantified the similarity between neural RDMs and model RDMs with regression or correlation.

#### High-vs-low dimensional + Train-test stimuli separation

This analysis was performed on all stimuli in the treasure hunt task. We regressed the neural RDM against three model RDMs: feature, location and train-test separation. The feature RDM came from a high-dimensional representation where stimuli were transformed into two-hot encoding (colour and shape), then the model RDM was calculated using hamming distance. The location RDM came from a low-dimensional representation where stimulus dissimilarity corresponded to spatial proximity on the ground-truth treasure map. The train-test separation assigned a distance of zero to training-training stimulus pairs and test-test stimulus pairs, and a distance of 1 to training-test stimulus pairs.

#### High-vs-low dimensional (feature vs location) representation of test stimuli

For this analysis, we regressed the neural RDM of test stimulus representations on two model RDMs: feature and location. The two model RDMs were specified in the same way as above.

#### Spatial representation of pirate positions

For this analysis, we computed the correlation between the neural RDM of pirate positions in the localiser task and a location RDM computed from the spatial proximities of the pirate positions.

#### Cross-task RSA

For this analysis, we constructed a cross-task neural RDM with entries reflecting the neural distance between a training stimulus and a pirate position. Since each pirate position corresponds to one of the training stimulus locations, we first tested whether matching stimulus-position pairs (diagonal entries in the neural RDM) were represented more similarly than non-matching pairs (off-diagonal entries). We computed the difference between the mean neural distance of the matching pairs and the mean neural distance of the non-matching pairs.

To examine whether the dissimilarity of non-matching pairs was better explained by a high-dimensional or low-dimensional model, we first symmetrized the asymmetric cross-task RDMs by averaging across the diagonal. We then regressed the lower triangle of each symmetric RDM onto the model RDMs. The high-dimensional model predicts a difference between matching and non-matching pair, but no difference among non-matching pairs, and the low-dimensional model predicts neural distance modulated by spatial proximity.

### Visualization of neural representation using MDS

We visualized the neural representations using multi-dimensional scaling. To this end, we first rescaled each generalizer’s neural RDM to a range between 0 and 1. Then, we computed a group average RDM from the rescaled RDMs of all generalizers. Finally, we applied classical MDS to this group-average RDM, extracting three components using MATLAB’s cmdscale function.

### Group-level statistical tests for ROI analyses and whole-brain searchlight analyses

Group-level statistics in ROIs were conducted with permutation test (10000 shuffles) unless otherwise specified. For one-sample tests on the means, the sign of data was randomly flipped to generate the null. For paired sample tests on the means, data within a pair was randomly exchanged to generate the null. For correlation tests, the null was generated by randomly permuting the data of one variable while the data of the other variable remained unchanged.

In whole-brain searchlight RSA analyses, group-level T maps were generated using the second level analysis of one sample t-test in SPM12. We reported the clusters that satisfied either of the following criteria: 1) cluster that passed an FWE correction of P=0.05; 2) cluster with a peak that passed an FWE correction of P=0.05.

## Data and code availability

Code to replicate the human experiments (https://github.com/ZiluLiang/Pirate-fMRI_Task) and analyses (https://github.com/ZiluLiang/Pirate-fMRI_Analysis) is available on GitHub. In line with local ethics guidelines, the fMRI data extracted from the regions of interest and pre-processed for the multivariate analyses are available on OSF (https://doi.org/10.17605/OSF.IO/F6HRC).

## Acknowledgements

This research is supported by a Wellcome Trust Discovery Award (https://wellcome.org/grant-funding/schemes/discovery-awards) (227928/Z/23/Z) to C.S.. M.K.F. was funded by a Wellcome Trust Henry Dale Fellowship (223263/Z/21/Z), a UKRI-converted ERC Starting Grant (EP/X021815/1) and a Leverhulme Award in Psychology. This research was funded in whole, or in part, by the Wellcome Trust (203139/Z/16/Z, 203139/A/16/Z, 223263/Z/21/Z, 221794/Z/20/Z and WT100973AIA). L.G. was supported by funding from the Medical Research Council (UKRI-MRC) (MR/W01971X/1 and MR/W008939/1 awarded to Professor Jill X O’Reilly and Associate Professor Helen Barron respectively). For the purpose of open access, the author has applied a CC BY public copyright licence to any Author Accepted Manuscript version arising from this submission. We would like to thank María Ruz Cámara, Sophie Arana, Eleanor Holton, Jirko Rubruck, and Kai Sandbrink and Jacques Pesnot-Lerousseau for their assistance with fMRI data collection.

