## SupplementaryMaterial for "Distinct roles of hippocampus and neocortex in symbolic compositional generalization"

### Supplementary Figures and Tables

Figure S1 (linked to Figure 1).

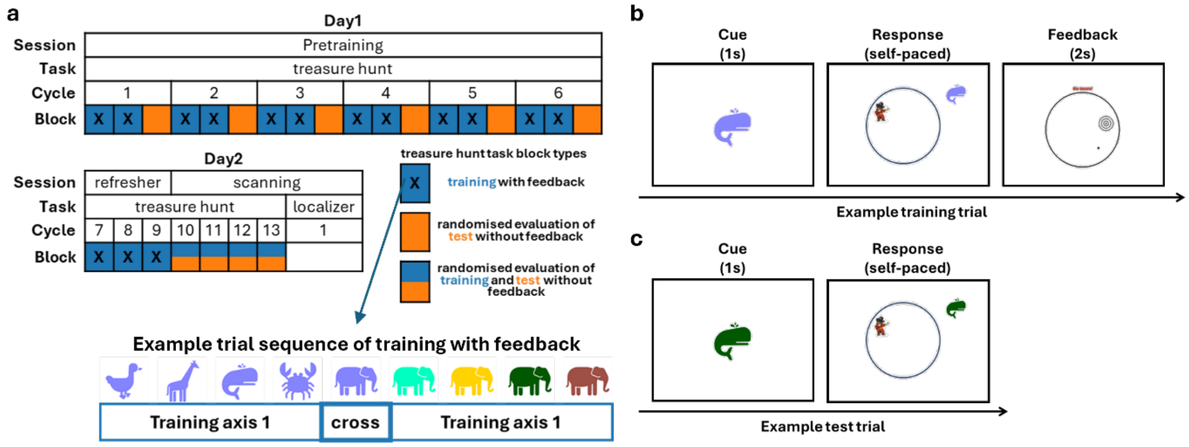


Figure S1. Experiment procedure and task timeline. (a) Procedure of the two-day experiment. (b-c) Trial timelines of the treasure hunt task in Day1. (b) Feedback was provided during training with feedback. (c) No feedback was provided during test.

Figure S2 (linked to Figure 1).

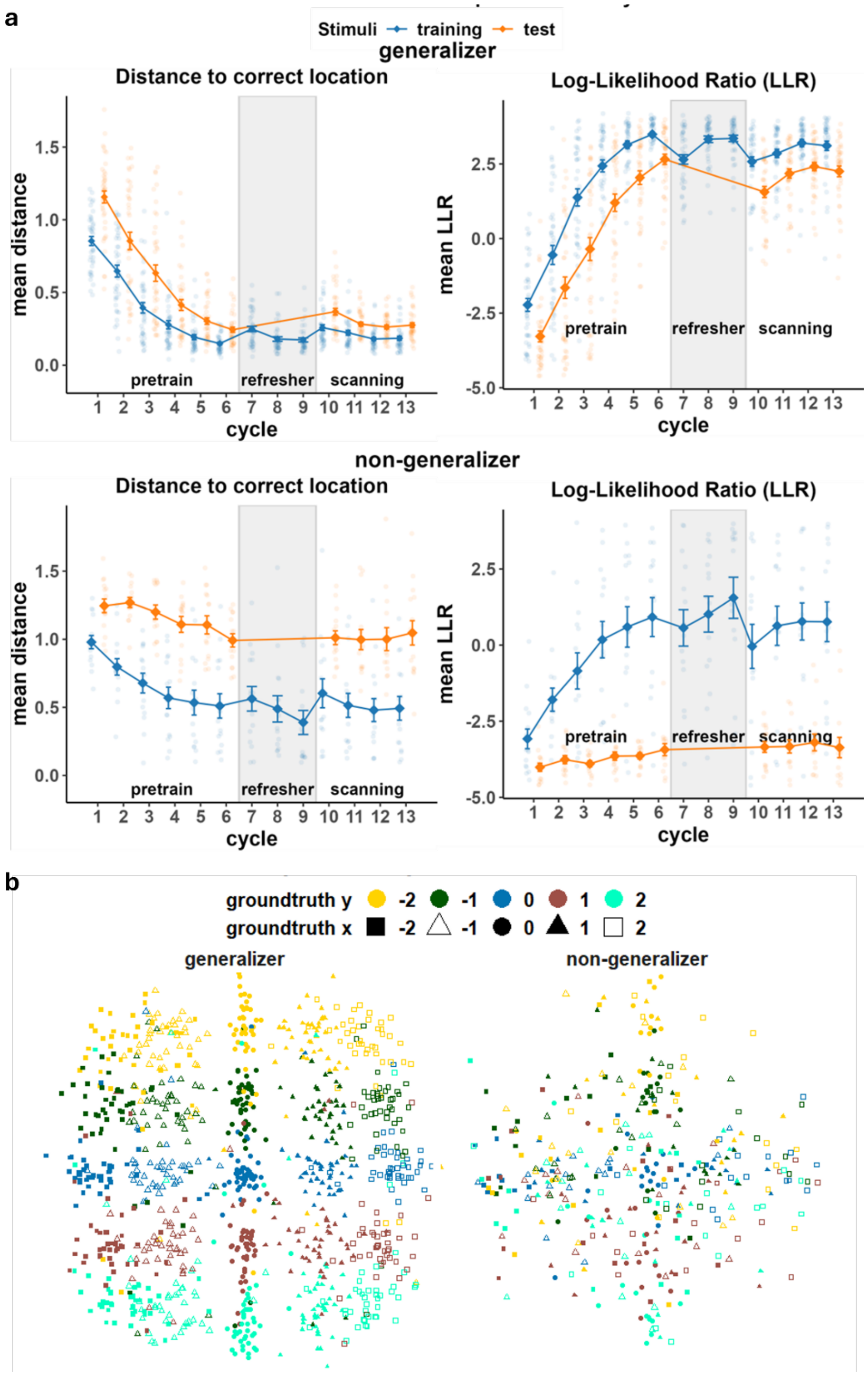


Figure S2. Behavioural performance. (a) Performance of generalizers (top panels) and non-generalizers (bottom panels) measured by distance to correct treasure location (left panels) and log-likelihood ratio (right panels). Translucent dots represent individual participants. Error bars represent mean ± standard error. (b) Participants' response in the scanner. Each dot represents one participant's response to a given stimulus averaged across all four runs of scanning.

Figure S3 (linked to Figure 2, Figure 5, and Figure 6).

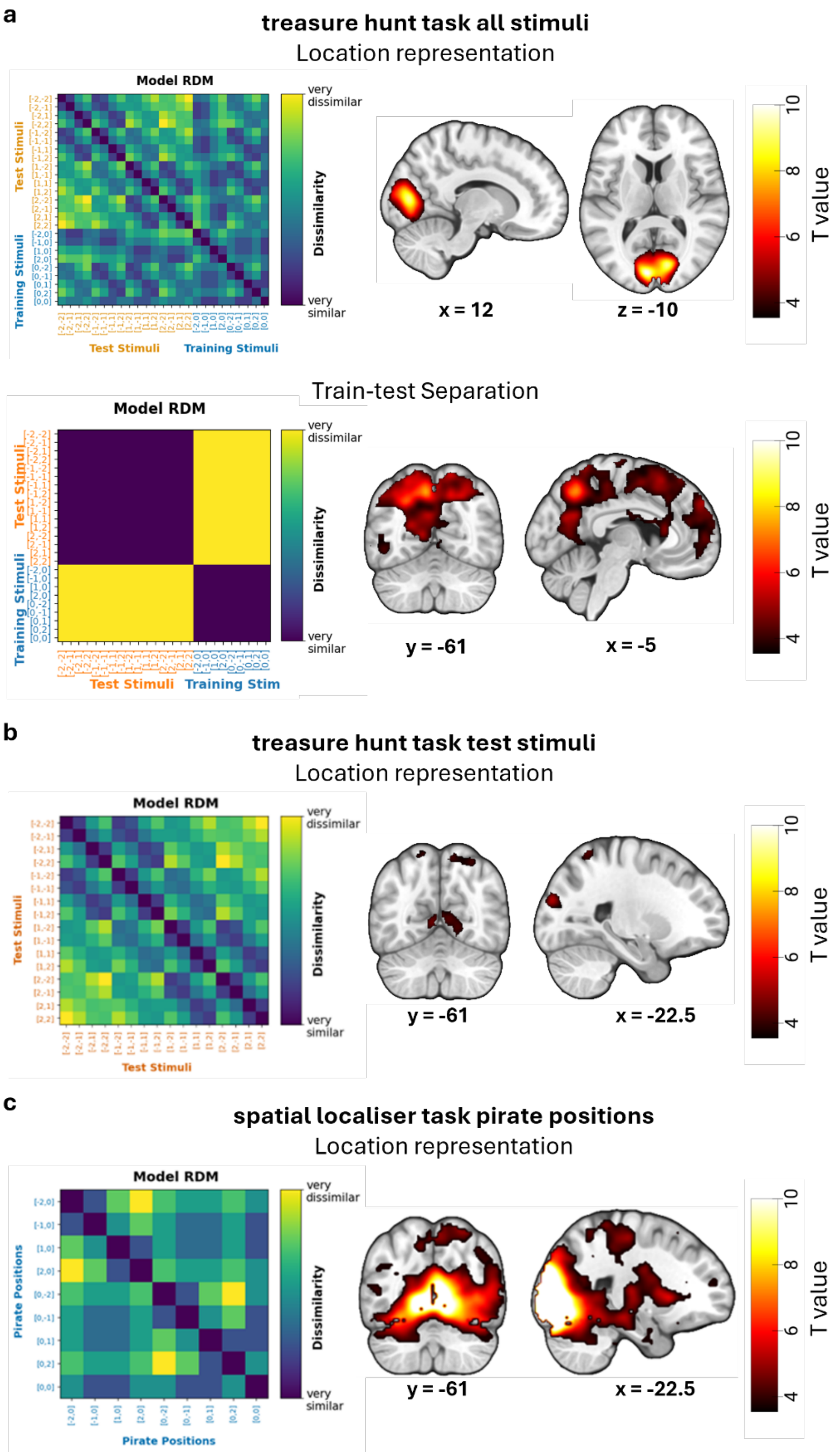


Figure S3. Whole-brain searchlight results. Panels on the left show the model RDMs, panels on the right show cluster corrected results. (a) Top: spatial location representation of all the stimuli in treasure hunt task. Bottom: separation between training stimuli and test stimuli. See Table S3 for the full table of searchlight results. Cluster-defining threshold was set at $p<0.0001$ to avoid oversized clusters. (b) Spatial location representation of only the test stimuli in treasure hunt task. See Table S4 for the full table of searchlight results. Cluster-defining threshold at $p<0.001$. (c) Spatial location representation of the pirate positions in the localiser task. See Table S5 for the full table of searchlight results. Cluster-defining threshold was set at $p<0.0001$ to avoid oversized clusters.

Figure S4 (linked to Figure 5).

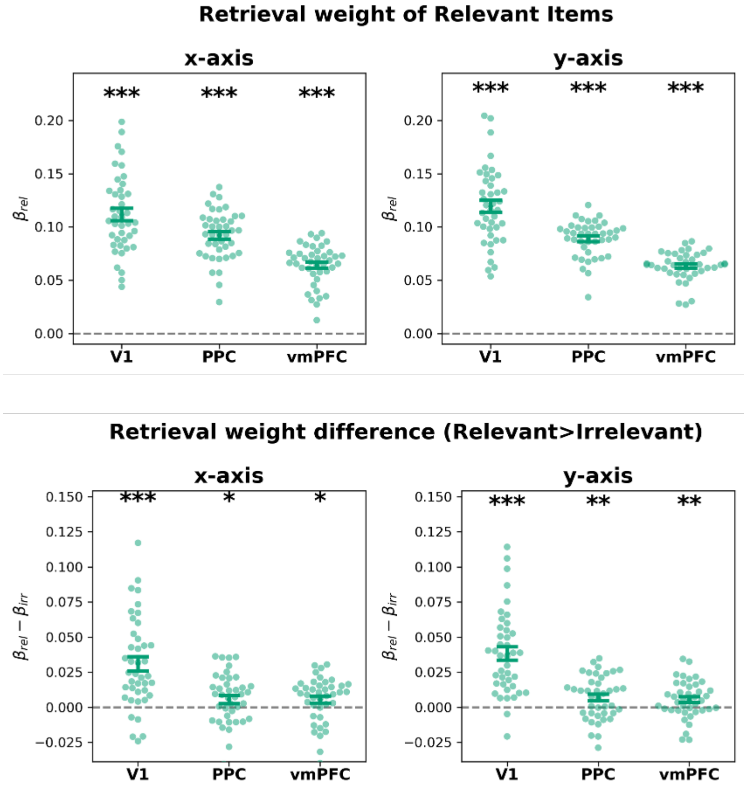


Figure S4. Sanity checks of retrieval pattern analysis. (a) Average weights of relevant ($\beta_{rel}$) and irrelevant training stimuli ($\beta_{irr}$) in the retrieval patterns for the three ROIs that showed vector addition composition. (b) Difference between weights of relevant and irrelvant training stimuli. Notes: Asterisks indicated results of permutation tests, *p<0.05, **p<0.01, ***p<0.001. Translucent dots represent the data from individual generalizers. Error bars represent mean ± standard error.

Figure S5 (linked to Figure 6).

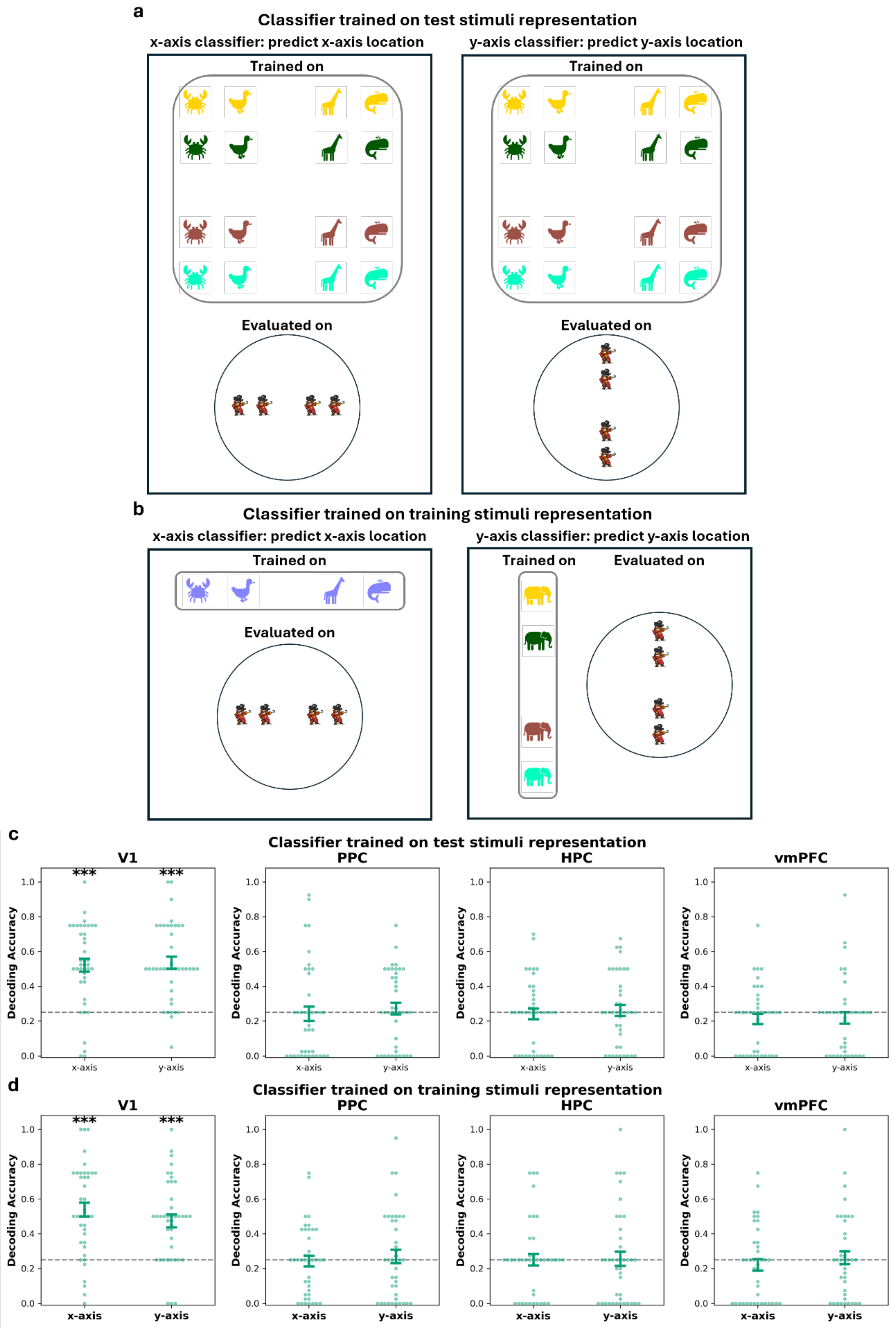


Figure S5. Cross-task decoding. (a-b) Schematic illustration of cross-task decoding procedure. First, classifiers were trained to predict the locations on a given axis based on the activity patterns of coloured animals from the treasure hunt task. Next, classifiers were evaluated on the data from the localiser task by applying it to predict pirate positions on a given axis based on the activity patterns of pirates from the localiser task. (a) Logistic regression classifiers were trained on the activity patterns of the test stimuli from the treasure hunt task to predict x-axis locations (left panel) or y-axis locations (right panel) of the test stimuli. (b) Same as (a) but the classifiers were trained on the activity patterns of the training stimuli. Note that when training classifiers to predict x-axis locations, only the activity patterns of the four training stimuli on x-axis were used. Similarly, when training classifiers to predict y-axis locations, only the activity patterns of the four training stimuli on y-axis were used. (c-d) Decoding accuracy in all four ROIs. (c) Classifiers were trained on test stimuli representation. (d) Classifiers were trained on training stimuli representation. Translucent dots represent the decoder's mean evaluation accuracy across 20 random seeds for each generalizer. Error bars represent mean ± standard error. Dashed grey lines represent chance level.

Table S1. Sanity checks for dimensionality estimates (linked to Figure 2).

a. Permutation test of reconstruction correlation in cross-validated SVD in generalizers.

| ROI | axis | mean correlation | p |
|---|---|---|---|
| V1 | x-axis | 0.151 | <0.001 |
| | y-axis | 0.157 | <0.001 |
| PPC | x-axis | 0.112 | <0.001 |
| | y-axis | 0.114 | <0.001 |
| HPC | x-axis | 0.031 | <0.001 |
| | y-axis | 0.030 | <0.001 |
| vmPFC | x-axis | 0.029 | <0.001 |
| | y-axis | 0.029 | <0.001 |

b. difference between dimensionality estimates of different training axes

| ROI | mean difference (x-axis > y-axis) | p |
|---|---|---|
| V1 | 0.073 | 0.288 |
| PPC | -0.104 | 0.100 |
| HPC | 0.061 | 0.707 |
| vmPFC | 0.024 | 0.880 |

c. correlation between dimensionality estimates of different training axes

| ROI | correlation | p |
|---|---|---|
| V1 | 0.593 | 0.016 |
| PPC | 0.588 | 0.054 |
| HPC | 0.363 | 0.027 |
| vmPFC | 0.216 | 0.192 |

Table S2. Linear mixed-effect models on correspondence between SVD projection and stimulus axis locations (linked to Figure 2).

(a) model summary

| | V1 | PPC | HPC | vmPFC |
|---|---|---|---|---|
| (Intercept) | 0.000 | 0.000 | 0.000 | 0.000 |
| | [-7.759, 7.759] | [-6.057, 6.057] | [-2.974, 2.974] | [-7.187, 7.187] |
| Axis Location | 33.331 *** | 15.474 *** | 7.031 *** | 17.001 *** |
| | [23.516, 43.145] | [7.812, 23.135] | [3.269, 10.794] | [7.911, 26.092] |
| Training Axis | 0.000 | 0.000 | 0.000 | 0.000 |
| | [-10.973, 10.973] | [-8.566, 8.566] | [-4.206, 4.206] | [-10.164, 10.164] |
| Axis Location× Training Axis | 6.088 | 11.526 * | 0.150 | 14.312 * |
| | [-7.792, 19.968] | [0.691, 22.361] | [-5.171, 5.471] | [1.456, 27.168] |
| N | 328 | 328 | 328 | 328 |
| N (participant) | 41 | 41 | 41 | 41 |
| AIC | 3494.889 | 3334.413 | 2873.585 | 3445.241 |
| BIC | 3517.647 | 3357.171 | 2896.343 | 3467.999 |
| R2 (fixed) | 0.245 | 0.162 | 0.077 | 0.153 |
| R2 (total) | 0.245 | 0.162 | 0.077 | 0.153 |

*** $p < 0.001$; ** $p < 0.01$; * $p < 0.05$.

(b) Post-hoc test on slopes of Axis Location

| ROI | Training Axis | slope | 95%CI | T | p |
|---|---|---|---|---|---|
| V1 | x-axis | 33.331 | [23.474, 43.187] | 6.656 | 1.45E-10 |
| | y-axis | 39.418 | [29.562, 49.275] | 7.872 | 7.41E-14 |
| PPC | x-axis | 15.474 | [7.779, 23.168] | 3.958 | 9.54E-05 |
| | y-axis | 26.999 | [19.305, 34.694] | 6.907 | 3.25E-11 |
| HPC | x-axis | 7.031 | [3.253, 10.810] | 3.663 | 0.000297 |
| | y-axis | 7.182 | [3.403, 10.960] | 3.741 | 0.000222 |
| vmPFC | x-axis | 17.001 | [7.872, 26.131] | 3.666 | 0.000294 |
| | y-axis | 31.314 | [22.184, 40.443] | 6.751 | 8.24E-11 |

Table S3. Searchlight RSA results: regressing all stimuli neural RDM on three model RDMs: feature RDM, location RDM, train-test separation RDM (linked to Figure 2 and Figure 5).

a. location RDM

| Anatomical Label | Peak coordinates (MNI) | | | Peak t value | Peak $p_{FWE}$ | Cluster | |
|---|---|---|---|---|---|---|---|
| | x | y | z | | | Size | $p_{FWE}$ |
| Calcarine_L | -5.0 | -86.0 | 10.5 | 10.193 | <0.001 | 1948 | <0.001 |
| Calcarine_R | 12.5 | -78.5 | 8.0 | 9.465 | <0.001 | 1948 | |
| Precentral_L | -40.0 | -18.5 | 58.0 | 5.483 | 0.012 | 96 | 0.023 |

b. train-test separation RDM

| Anatomical Label | Peak coordinates (MNI) | | | Peak t value | Peak $p_{FWE}$ | Cluster | |
|---|---|---|---|---|---|---|---|
| | x | y | z | | | Size | $p_{FWE}$ |
| Parietal_Inf_L | -35.0 | -51.0 | 40.5 | 7.847 | <0.001 | 4848 | 0.0000 |
| Precuneus_L | -7.5 | -58.5 | 45.5 | 7.661 | <0.001 | | |
| Parietal_Inf_L | -25.0 | -58.5 | 40.5 | 6.950 | <0.001 | | |
| Frontal_Mid_2_L | -37.5 | 46.5 | -9.5 | 7.313 | <0.001 | 5129 | 0.0000 |
| Frontal_Mid_2_L | -37.5 | 34.0 | 33.0 | 7.311 | <0.001 | | |
| Frontal_Inf_Tri_L | -42.5 | 34.0 | 18.0 | 7.187 | <0.001 | | |
| Supp_Motor_Area_R | 12.50 | -13.50 | 58.00 | 6.10 | <0.001 | 175 | 0.0015 |
| Precentral_R | 27.50 | -6.00 | 53.00 | 5.43 | 0.022 | | |
| Frontal_Sup_2_R | 22.50 | -3.50 | 63.00 | 4.49 | 0.232 | | |
| Frontal_Mid_2_R | 35.00 | 51.50 | 15.50 | 5.88 | 0.007 | 225 | 0.0005 |
| Frontal_Mid_2_R | 42.50 | 44.00 | 13.00 | 5.06 | 0.059 | | |
| Hippocampus_R | 37.50 | -11.00 | -14.50 | 5.73 | 0.010 | 37 | 0.0752 |
| Brain stem | 2.50 | -31.00 | -29.50 | 5.27 | 0.034 | 28 | 0.1071 |
| Paracentral_Lobule_L | -15.00 | -26.00 | 73.00 | 4.91 | 0.086 | 50 | 0.0469 |
| Cingulate_Mid_R | 7.50 | -26.00 | 48.00 | 4.77 | 0.121 | 136 | 0.0038 |
| Paracentral_Lobule_L | -7.50 | -31.00 | 53.00 | 4.51 | 0.220 | | |
| Cingulate_Mid_L | -7.50 | -33.50 | 43.00 | 4.35 | 0.316 | | |
| white matter | 20.00 | -21.00 | 33.00 | 4.77 | 0.122 | 108 | 0.0080 |
| white matter | 27.50 | -23.50 | 20.50 | 4.47 | 0.242 | | |
| Insula_R | 37.50 | -11.00 | 18.00 | 4.45 | 0.253 | | |
| Insula_L | -42.50 | -1.00 | 0.50 | 4.76 | 0.124 | 35 | 0.0812 |
| Central Opercular Cortex | -45.00 | 1.50 | -7.00 | 4.16 | 0.445 | | |
| Frontal_Sup_Medial_R | 15.00 | 49.00 | 0.50 | 4.72 | 0.138 | 57 | 0.0370 |
| ACC_pre_R | 15.00 | 39.00 | 0.50 | 4.54 | 0.210 | | |
| Frontal_Sup_2_R | 17.5 | 59.0 | 3.0 | 4.346 | 0.316 | | |

Note:

[1]The reported second level results for these two model RDM employed a more stringent cluster-defining threshold at p<0.0001 to avoid oversized clusters.

[2]Anatomical labels were from AAL. When AAL labels were not available, the Harvard-Oxford cortical and subcortical atlas was used.

Table S4. Searchlight RSA results: regressing test stimuli neural RDM on two model RDMs: feature RDM and location RDM (linked to Figure 2 and Figure 5).

| Anatomical Label | Peak coordinates (MNI) | | | Peak t value | Peak $p_{FWE}$ | Cluster | |
|---|---|---|---|---|---|---|---|
| | x | y | z | | | Size | $p_{FWE}$ |
| C mialcarine_R | 12.5 | -78.5 | 13.0 | 8.526 | <0.001 | 23932 | <0.001 |
| Calcarine_L | -7.5 | -86.0 | 10.5 | 8.114 | <0.001 | | |
| Cuneus_R | 15.0 | -88.5 | 35.5 | 3.472 | 0.905 | | |
| Precentral_L | -37.5 | -23.5 | 63.0 | 6.071 | 0.003 | 111 | 0.186 |
| Postcentral_L | -37.5 | -36.0 | 63.0 | 4.740 | 0.111 | 428 | 0.003 |
| Parietal_Inf_L | -57.5 | -23.5 | 48.0 | 4.047 | 0.479 | | |
| Temporal_Pole_Mid_L | -30.0 | 11.5 | -34.5 | 5.068 | 0.049 | 32 | 0.656 |
| Temporal_Pole_Mid_R | 35.0 | 19.0 | -34.5 | 4.896 | 0.075 | 39 | 0.598 |
| Precuneus_R | 12.5 | -66.0 | 55.5 | 4.071 | 0.460 | 225 | 0.051 |
| Parietal_Sup_R | 27.5 | -61.0 | 55.5 | 3.923 | 0.581 | | |
| Parietal_Sup_L | -22.5 | -56.0 | 65.5 | 4.245 | 0.333 | 102 | 0.249 |

Note:

[1]The reported second level result employed a cluster-defining threshold at p<0.001.

[2]We also reported some subthreshold clusters in grey if their contralateral cluster passed cluster correction or voxel-wise correction.

[3]Anatomical labels were from AAL. When AAL labels were not available, the Harvard-Oxford cortical and subcortical atlas was used.

Table S5. Searchlight RSA results: correlation between pirate position neural RDM and location model RDM (linked to Figure 6).

| Anatomical Label | Peak coordinates (MNI) | | | Peak t value | Peak $p_{FWE}$ | Cluster | |
|---|---|---|---|---|---|---|---|
| | x | y | z | | | Size | $p_{FWE}$ |
| Lingual_R | 5.0 | -81.0 | -2.0 | 33.555 | <0.001 | 21010 | <0.001 |
| Lingual_L | -7.5 | -83.5 | -2.0 | 29.789 | <0.001 | 21010 | |
| Lingual_R | 20.0 | -86.0 | -7.0 | 26.481 | <0.001 | 21010 | |
| Location not in atlas | 37.5 | -6.0 | -14.5 | 5.477 | 0.010 | 213 | 0.006 |
| Fusiform_R | 35.0 | -6.0 | -34.5 | 4.454 | 0.133 | 213 | |
| ParaHippocampal_R | 25.0 | -1.0 | -34.5 | 4.349 | 0.169 | 213 | |
| OFCpost_L | -45.0 | 24.0 | -14.5 | 4.977 | 0.036 | 76 | 0.043 |
| Frontal_Mid_2_L | -42.5 | 24.0 | 43.0 | 4.857 | 0.049 | 52 | 0.067 |

Note:

[1]The reported second level result employed a more stringent cluster-defining threshold at $p<0.0001$.

[2]Anatomical labels were from AAL. When AAL labels were not available, the Harvard-Oxford cortical and subcortical atlas was used.
